# Distinct mechanisms of antibody-mediated HCV neutralization revealed by nanobody-guided epitope mapping

**DOI:** 10.64898/2026.08.01.742232

**Authors:** Haneen Tarabih, Jenna Weisz, Itai Yechezkel, Els Pardon, Nele Buys, Jan Steyaert, Erick Giang, Rina Fraenkel, Petr Pompach, Mansun Law, Netanel Tzarum

## Abstract

Hepatitis C virus (HCV) remains a major global health challenge despite the availability of highly effective antiviral therapies, underscoring the need for a broadly protective vaccine. The envelope glycoprotein E2 is the principal target of neutralizing antibodies, yet the full repertoire of vulnerable epitopes and mechanisms of antibody-mediated neutralization remains incompletely understood. Here, we exploited the unique binding properties of camelid nanobodies to probe the antigenic landscape of HCV E2 beyond the immunodominant human antibody response. We isolated a diverse panel of E2-specific nanobodies, including broadly neutralizing antibodies with high-affinity cross-reactivity toward genetically diverse HCV isolates. By combining cross-neutralization assays, competition binding experiments, and high-resolution hydrogen–deuterium exchange mass spectrometry (HDX-MS), we identified three mechanistically distinct classes of neutralizing epitopes. While one class targets the canonical E2 neutralization face, a second class recognizes antigenic region 1 (AR1), independently validating and extending recent evidence that this region represents a functional site of viral vulnerability. These findings demonstrate that broadly neutralizing antibody responses extend beyond the canonical neutralization face and establish a broader framework for understanding HCV neutralization. More broadly, our study illustrates how alternative antibody repertoires can reveal functionally important antigenic surfaces that are underrepresented in conventional human antibody responses, providing new opportunities for the rational design of next-generation HCV vaccine immunogens.

## Introduction

Hepatitis C virus (HCV) remains a major global health challenge despite the availability of highly effective direct-acting antiviral therapies ^2–5^. According to the World Health Organization’s latest report, an estimated 50 million people worldwide are chronically infected with HCV, with about 1.0 million new infections occurring per year (World Health Organization, Hepatitis C: Fact sheet (2025), https://www.who.int/news-room/fact-sheets/detail/hepatitis-c). Chronic infection is a leading cause of liver cirrhosis, hepatocellular carcinoma, and liver failure, resulting in hundreds of thousands of deaths annually. Furthermore, new HCV infections continue to occur, mostly in at-risk groups, such as people who inject drugs (PWIDs) ^6^, underscoring the persistent need for a preventive vaccine to reduce transmission and disease burden. Thus, a prophylactic vaccine that elicits potent and broadly neutralizing immune responses is critical for the eradication of HCV.

A major challenge in HCV vaccine development is the virus’s extreme genetic and antigenic diversity, including eight distinct genotypes (Gt) ^7^, as well as the immune evasion strategies employed by its envelope proteins (Env). Nevertheless, spontaneous viral clearance occurs in approximately 30% of acutely infected patients, indicating that chronic HCV infection is preventable if an effective immune response can be elicited by vaccination. HCV clearance is strongly associated with an early and broad neutralizing antibody (bnAb) response during the acute phase of infection ^8^.

HCV is an enveloped, positive-sense single-stranded RNA virus. Viral entry is facilitated by the Env glycoproteins E1 and E2, which form a noncovalent heterodimer on the surface of the virion. The E2 glycoprotein (amino acids 384–746) includes an N-terminal hypervariable region 1 (HVR1), a central core domain containing variable regions 2 and 3 (VR2 and VR3), and a C-terminal stalk that connects the ectodomain to the transmembrane domain (TM) (Supplementary Fig. 1A). The stalk region is wrapped and stabilized by E1, effectively connecting the E2 core domain to the TM and enhancing heterodimer stability, as shown in the E1E2 cryo-EM structure ^9^. E2 acts as the primary receptor-binding protein of HCV, interacting with multiple host entry factors, with CD81 and scavenger receptor class B type I (SR-BI) being the most crucial ^10^. Consequently, E2 is the primary target of nAbs, which aid in viral clearance and protective immunity.

High-resolution, molecular-level characterization of HCV Env neutralization epitopes is essential for designing cross-reactive vaccine antigens that induce high levels of bnAbs ^11,12^. Advances in structural studies of HCV Env over the past fifteen years have contributed to the characterization of E2 antigenic sites and to the understanding of how Abs recognize HCV^13–15^. The main and most studied antigenic surface of the HCV Env was defined as the neutralization face, a predominantly discontinuous hydrophobic surface comprising the front layer (FL) and the CD81 binding loop (CD81bl), which overlaps with the CD81 receptor binding site on E2 (Fig. 1A and Supplementary Fig. 1A). The neutralization face overlaps with three E2 main neutralization sites: antigenic site 412 (AS412), antigenic site 434 (AS434), and antigenic region 3 (AR3) ^16,17^. AR3 is a cluster of discontinuous epitopes that overlap the neutralization face (Fig. 1A) and is the target for numerous highly potent bnAbs.

**Fig.1.**
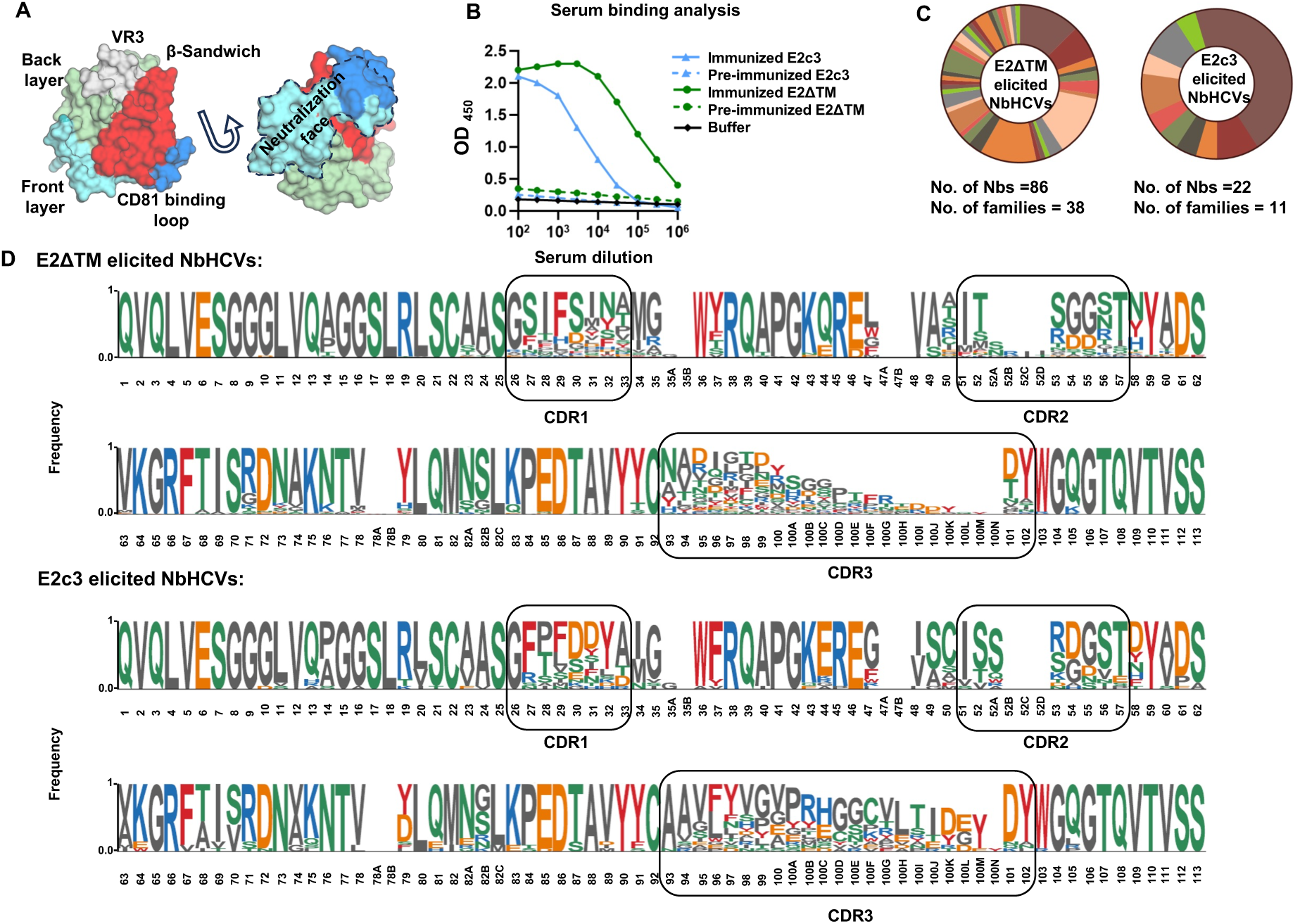
Generation and characterization of an E2-specific NbHCV library. (**A**) Structure of the HCV H77 E2 core (E2c; PDB: 4MWF) shown as a molecular surface colored by structural elements: the central immunoglobulin-like β-sandwich domain (red), the N-terminal front layer (cyan), and the C-terminal back layer (green). The extended CD81-binding loop is shown in blue. The neutralizing face of E2 is indicated by a dashed line. (**B**) ELISA binding of pre-immune and immune serum samples from llama immunization experiments to the purified E2 antigens, demonstrating robust induction of E2-specific HCAb responses. (**C**) Classification of E2-specific HCAbs into families based on heavy chain complementarity-determining region 3 (CDRH3) length and sequence similarity. (**D**) Sequence logo representation of the NbHCV library elicited by E2ΔTM (upper) and E2c3 (lower). Each NbHCV contains a unique CDRH3 sequence. The frequency of each amino acid at each position is indicated. FR, framework region.

AR3-targeting neutralization responses are dominated by bnAbs encoded by the IGHV1-69 (V_H_1-69) heavy-chain variable germline gene family ^13,18,19^. Structural studies of E2-AR3-targeting bnAb complexes indicated that they share similar binding features yet adopt distinct binding modes ^9,13,17,18,20–27^. Moreover, structural studies indicated that E2 FL can adopt two conformations upon binding to an AR3-targeting bnAb ^22^.

Recently, Ogega et al. performed longitudinal B cell receptor repertoire analysis in an individual who spontaneously cleared multiple HCV infections, identifying a large panel of genetically diverse bnAbs that utilize a broad range of VH gene segments ^23^. Structural characterization of two E2-bnAb complexes, encoded by the V_H_1-46 gene, revealed that they bind to AR3, whereas an nAb encoded by the V_H_4-34 gene binds to an antigenic site considered a non-neutralization site. More recently, a study investigating the B cell response in rhesus macaques (RM) vaccinated with the Chiron E1E2 vaccine indicated that the AR3 can be targeted by bnAbs encoded by germline genes equivalent to V_H_1-69 and V_H_4-59 ^24^. These findings suggest that effective vaccines should aim to elicit bnAbs against multiple E2 epitopes rather than focusing on a single dominant site, and that genetic diversity among induced bnAbs may be beneficial.

In this study, we aim to characterize new neutralization epitopes on the E2 protein using heavy-chain-only Abs (HCAbs), natural single-domain Abs found in Camelidae sera ^28^. HCAbs have proven particularly effective for probing viral Env proteins and for guiding vaccine immunogen design, owing to their small size, structural stability, and ability to access recessed or conformational epitopes that are often poorly targeted by conventional Abs ^29,30^. Note that neutralizing HCAbs against E2 have been reported ^31^, indicating the probability of eliciting E2-specific HCAbs. To identify new HCV neutralization epitopes, camelids were immunized with E2 antigens. RNA encoding the variable domains of HCAbs was isolated from lymphocytes to construct a phage-display library of E2-specific Nbs. From this library, we selected, expressed, and purified 64 E2-specific Nbs (NbHCVs), 24 of which cross-bound E1E2 antigens from multiple genotypes. Neutralization and binding-competition assays indicated that broad neutralization is primarily mediated by blocking the interaction between E2 and the host receptor CD81. Epitope mapping experiments further classified the NbHCVs into three distinct groups. Class 2 NbHCVs target the well-defined E2 neutralization face, while class 1 NbHCVs bind an antigenic region previously considered non-neutralizing that has recently emerged as an additional neutralization site. Overall, these findings expand the known range of HCV neutralizing epitopes and offer new insights to guide the design of improved E2 immunogens for next-generation HCV vaccines.

## Results

### Camelid immunization with soluble E2 elicits robust E2-specific HCAb responses

To identify and characterize novel neutralization epitopes, camelids were immunized with E2 envelope proteins to isolate E2-specific HCAbs. To maximize the diversity of elicited HCAbs, two antigens were used: the soluble E2 ectodomain from the H77 prototypic strain (H77 E2ΔTM; amino acids 384–645) and the H77 E2c3 core domain (E2 residues 412–645 with an internal truncation of variable region 2 and 3 and removal of the N448 and N576 glycosylation sites; Supplementary Fig.1A) ^21^. E2ΔTM contains HVR1, which modulates interactions with the SR-BI receptor, as well as the stalk region that interacts with E1. Immunization with E2ΔTM may therefore elicit HCAbs that target new neutralization epitopes, e.g., by blocking the SR-BI binding site or by preventing conformational changes necessary for HCV entry. In contrast, E2c3 lacks the flexible N- and C-terminal regions and the variable domains of E2, which can interfere with the induction of neutralizing responses. Consequently, E2c3 was expected to preferentially elicit higher titers of HCAbs targeting the conserved neutralization face of E2 (Fig. 1A).

Two llamas, each immunized with a different antigen, received six consecutive injections of highly purified E2s, and blood was collected three days after the final immunization (Supplementary Fig. 1B). A serum-binding assay showed high levels of E2-specific Abs, with a stronger binding response in the E2ΔTM-immunized sample (the assay measures total Ab response, including HCAbs and conventional Abs; Fig. 1B). Total RNA was extracted from peripheral blood lymphocytes, and cDNA was synthesized. The HCAbs variable regions (named nanobodies, Nbs) were amplified and cloned to generate two independent E2-specific phage-display libraries. Panning experiments on these libraries demonstrated enrichment of E2-specific Nbs (Supplementary Table 1). Using the enriched sub-libraries, we selected individual E2-specific clones, screened them in ELISA, and then sequenced positive clones to produce the final HCV-specific Nb (NbHCVs) library. The library, derived from both immunization experiments, contains 106 distinct E2-specific clones, 86 from the E2ΔTM immunization and 20 from the E2c3 immunization (Fig. 1C). All clones are grouped into 48 NbHCV families based on the length and sequence similarity of the complementarity-determining region heavy chain 3 (CDRH3, same length and over 80% sequence identity) (Fig. 1C) ^32^. Note, Nbs from the same family are usually derived from the same B-cell lineage and bind to the same epitope on the target ^32^.

Sequence analysis of the NbHCV libraries showed high sequence diversity, indicated by a wide range of CDRH3 lengths (5-23 amino acids for the E2ΔTM-derived library and 10-24 amino acids for the E2c3-derived library). A sequence logo displayed conserved framework residues, with significant variation in the CDRH loops (Fig. 1D). Phylogenetic analysis further supported the CDRH3-based family classification, as members of the same family generally clustered on neighboring or closely related branches rather than forming strictly separate clades (Supplementary Fig. 1C).

### NbHCVs showed high affinity and cross-binding to HCV genotypes

To evaluate the binding specificity of the NbHCVs, representative Nbs from each family were produced as recombinant proteins. NbHCVs were chosen based on two criteria: including at least one Nb from each family and preferential selection of clones showing higher binding signals during the antigen-binding step (data not shown). For families with many members (e.g., families 1 and 13), multiple representative Nbs were expressed. In total, 64 E2-specific NbHCVs (53 from the E2ΔTM library and 11 from the E2c3 library) were cloned as His-tagged fusion proteins into the pMESy4 expression vector for periplasmic production ^32^, expressed in the E. coli WK6 strain, and purified through affinity and size-exclusion chromatography (SEC) (Supplementary Fig. 2A).

To evaluate binding specificity, recombinant NbHCVs were tested for their ability to bind native E1E2 heterodimers captured from transfected cell lysates, as well as soluble H77 E2ΔTM and E2c3 used for immunization, using an enzyme-linked immunosorbent assay (ELISA). Note that Nbs, unlike conventional IgGs, are monovalent binding domains and significantly smaller in size (∼15 kDa compared to ∼150 kDa). E2ΔTM-derived NbHCVs revealed three distinct binding patterns (Fig. 2A). A subset of NbHCVs bound all three antigens, indicating recognition of epitopes within the E2c3 core that remain accessible in the native E1E2 complex (e.g., NbHCV089, NbHCV091, and NbHCV135). The second subset bound soluble E2ΔTM and E2c3 but showed reduced binding to native E1E2, suggesting recognition of epitopes that are less accessible in the native E1E2 complex. The third subset exhibited low binding across all tested antigens. NbHCVs derived from the E2c3 library generally showed weaker binding and were not pursued further (Supplementary Fig. 2B). Based on binding signal strength and assay reproducibility, 24 E2ΔTM-derived NbHCVs representing 14 families were selected for further analysis (24 NbHCVs panel).

**Fig.2.**
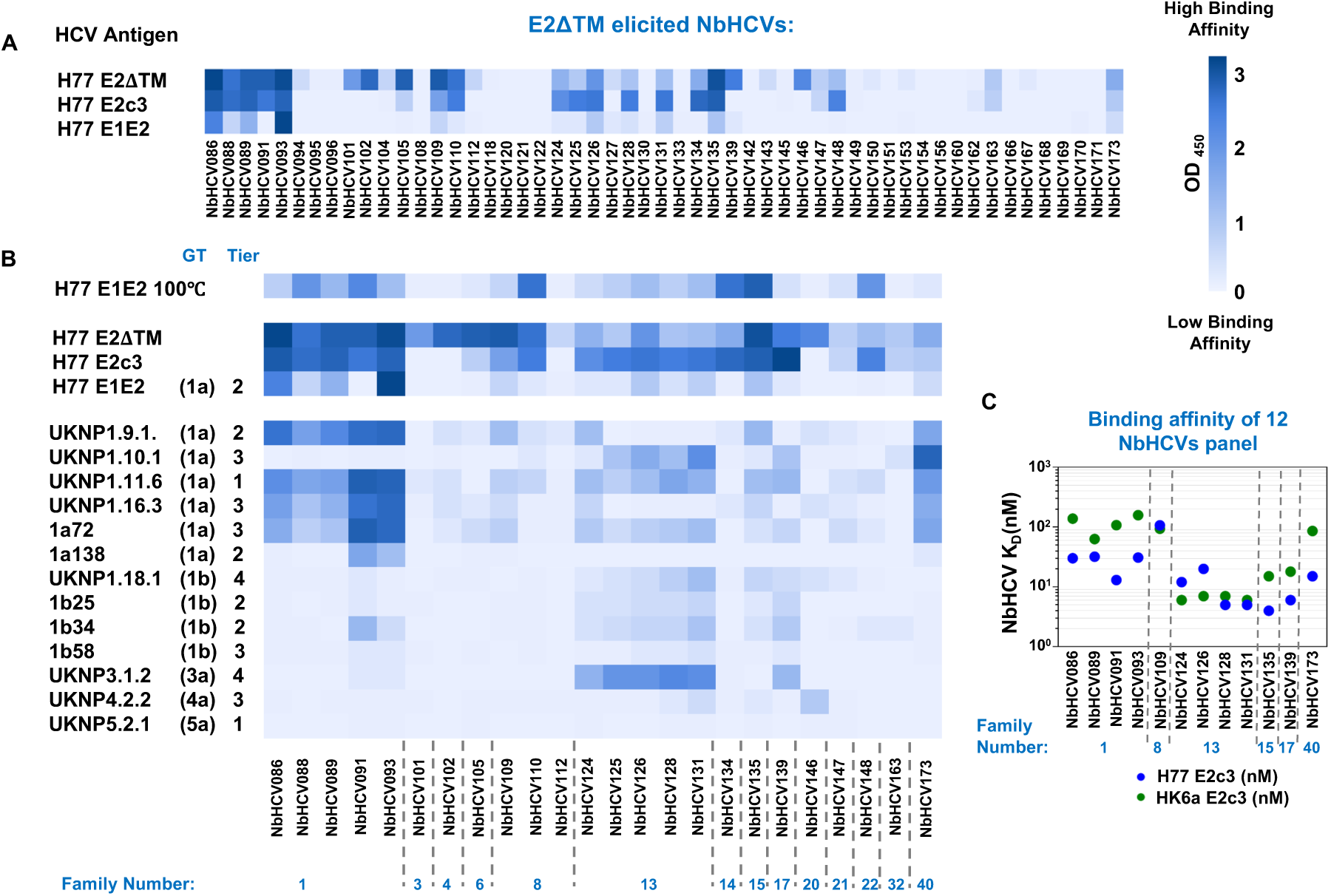
High-affinity cross-binding of NbHCVs to HCV envelope antigens across genotypes. (**A**) ELISA binding analysis of representative NbHCVs elicited by E2ΔTM immunization against autologous H77 antigens. Binding of NbHCVs to native H77 E1E2 and soluble E2ΔTM and E2c3 proteins is shown as a heat map of optical density values at 450 nm (OD450), with dark blue indicating stronger binding. Data represent mean OD values from duplicate wells within a single experiment. Results are representative of at least two independent experiments. (**B**) ELISA binding analysis of the 24 NbHCV panel against the native E1E2 HCV reference panel 1. The genotype and Tier of each isolate are indicated. (**C**) Binding affinity determination of the 12 NbHCV panel against H77 and HK6a E2c3 antigens. KD values were determined by bio-layer interferometry (BLI).

To test the NbHCVs cross-binding breadth, ELISA experiments were performed against the representative, antigenically diverse HCV reference panel reported by Salas et al. ^1^, which spans multiple tiers of neutralization sensitivity and has been proposed as a standard for measuring neutralizing breadth and potency of anti-HCV Abs across laboratories (Fig. 2B). In parallel, we used a representative Gt 1-6 panel used in our previous studies (Supplementary Fig. 2C) ^21,33^. The results using the Salas panel indicated a broad spectrum of cross-binding capabilities across the families. For instance, NbHCVs from families 1 and 40 show a preference for high binding of genotype 1a E1E2s, spanning from tiers 1-3, while NbHCVs from families 13, 15, and 17 show moderate binding to a broader part of the panel, including E1E2 from genotype 1a, 1b, and 3a.

To determine binding affinity, the equilibrium dissociation constants (K_D_) of the 24 NbHCV panel for H77 E2c3 were measured using biolayer interferometry (BLI) (Fig. 2C and Supplementary Fig. 2D, E). The NbHCVs exhibited high binding affinities, with K_D_ values in the nanomolar range (2–530 nM). Based on the ELISA results (Fig. 2B), a panel of 12 representative NbHCVs was selected for further analysis (the 12 NbHCV panel). The cross-binding affinity of these NbHCVs to the heterologous HK6a E2c3 was then evaluated. All NbHCVs displayed high binding affinity, although differences were observed between Nb families. Notably, NbHCVs from families 8 and 13 exhibited comparable or higher affinity for the heterologous E2c3 than for the autologous antigen, whereas NbHCVs from the remaining families showed stronger binding to the autologous protein.

### NbHCVs demonstrated cross-neutralizing activity

Neutralization of HCV pseudoparticles (HCVpp) by the 12-NbHCV panel was first tested against the autologous H77pp. NbHCVs from families 1, 8, and 13 exhibited potent activity, achieving >50% neutralization with IC_50_ values of 0.4-12.5 μg/mL (Fig. 3 and Supplementary Fig. 3). Assessment of neutralization breadth using the antigenically diverse HCV reference panel (n = 14) revealed that family 13 NbHCVs exhibited the strongest cross-neutralizing activity, neutralizing ≥50% of isolates across multiple Gts and tiers at both 20 μg/mL and 2 μg/mL (Fig. 3 and Supplementary Fig. 3). NbHCVs from families 1 and 8 demonstrated moderate breadth, neutralizing 3–6 isolates at 20 μg/mL. Conversely, NbHCV135, NbHCV139, and NbHCV173 showed very weak neutralization activity across the panel.

**Fig.3.**
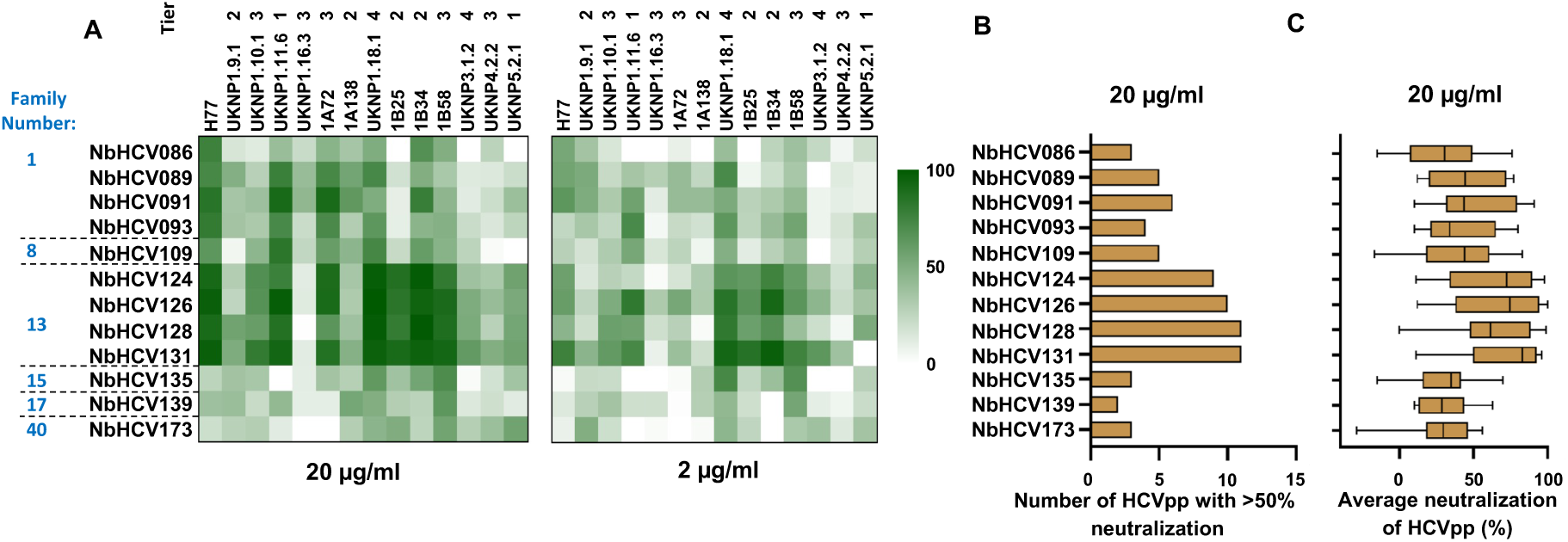
Cross-neutralization capability of the NbHCVs. (**A**) HCVpp neutralization assay against the Salas native E1E2 HCV reference panel (n=14). The percentage of neutralization at 20 µg/ml and 2 µg/ml is shown as a heat map, with dark green indicating 100% neutralization. (**B**) The number of HCVpp with >50% neutralization for each NbHCV at a concentration of 20 µg/ml. (**C**) The average neutralization of each NbHCV against the HCVpp panel at a concentration of 20 µg/ml. Lines, boxes, and whiskers indicate medians, 25%–75%, and 5%– 95% percentiles, respectively.

### Neutralizing NbHCVs target epitopes that overlap with the E2 neutralization face and Antigenic region 1

The binding and neutralization data showed that NbHCVs from different families have distinct binding profiles. To identify NbHCV epitopes, we performed competition ELISAs using domain-specific E2 Abs, including the bnAb AP33 targeting antigenic site 412 (AS412) ^34^, AR3A, and HC1AM targeting antigenic region 3 (AR3) ^21,22^, and a non-neutralizing Ab targeting antigenic region 1 (AR1) ^22^ (Fig. 4A). EC_75_ values were first determined for all NbHCVs (Supplementary Fig.4). Next, H77 E2c3 captured ELISA plates were pre-incubated with saturating concentrations of blocking Abs, followed by incubation with NbHCVs at their EC_75_ concentrations. In parallel, competition with the bnAb HEPC3 ^26^ and the HCV receptor CD81 was tested (for CD81, we used an Fc-fused large extracellular loop of CD81, CD81-LEL). EC_75_ value for HEPC3 and CD81-LEL were first determined (Supplementary Fig. 4). Next, H77 E2c3 captured ELISA plates were pre-incubated with saturating concentrations of the NbHCVs, followed by incubation with HEPC3 and CD81 LEL at their EC_75_ concentration. The competition assays showed that NbHCVs from families 13 and 40 strongly compete with AR3-specific antibodies (AR3A, HC1AM, and HEPC3) and CD81-LEL (Fig. 4A, B). NbHCVs from families 1 and 8 strongly compete with AR3A and HC1AM but exhibit only moderate competition with HEPC3 and CD81-LEL. In contrast, NbHCV135 and NbHCV139 show no competition with AR3-specific antibodies or CD81-LEL. No inhibition was observed with the AS412-specific antibody AP33. Notably, NbHCV binding was also inhibited by the AR1-targeting Ab (E1), that target an epitope located at the C-terminal part of the β-sandwich region (Supplementary Fig.1A), which is spatially distinct from the E2 neutralization face and has recently been reported as an epitope for nAbs ^23^.

**Fig.4.**
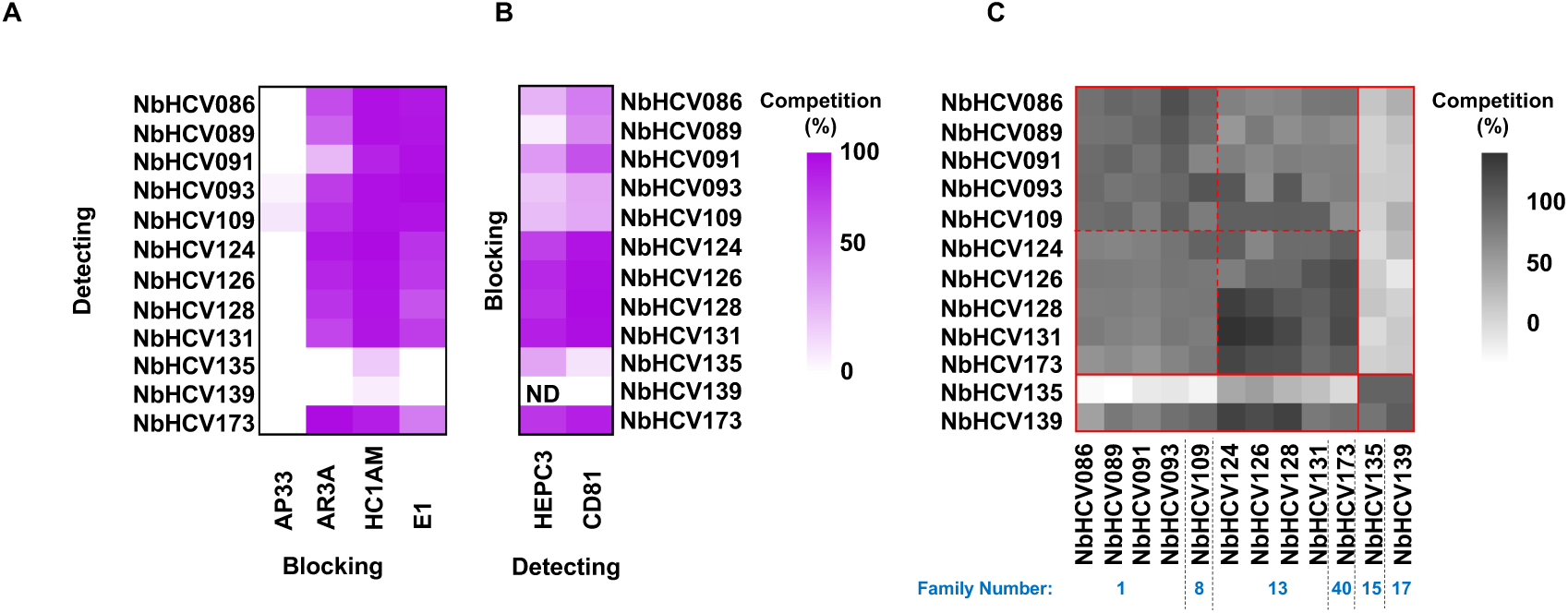
Epitope mapping of NbHCVs using competition binding assays. **(A)** Competitive ELISA assessing NbHCV binding in the presence of domain-specific monoclonal IgGs. IgG E1 is a non-neutralizing Ab that targets AR1. Domain-specific IgGs were used as competitors at saturating concentrations, and binding of His-tagged NbHCVs was detected at their EC_75_ levels. Percent inhibition was calculated by comparing the binding signal in the presence of a competitor with that in its absence, expressed as a percentage reduction relative to the no-competitor condition. Results are shown as a heat map of optical density at 450 nm (OD_450_), with dark purple indicating stronger inhibition. Data represent mean OD_450_ values from duplicate wells within a single experiment and are representative of two independent experiments. **(B)** Competitive ELISA assessing inhibition of HEPC3 and Fc-CD81-LEL binding by NbHCVs. NbHCVs were used as competitors at saturating concentrations, and HEPC3 or Fc-CD81-LEL binding was measured at the EC_75_. Percent inhibition was calculated as described in panel (A). ND, not detected. **(C)** BLI-based competitive epitope binding using the 12-NbHCV panel. Results are shown as a heat map indicating the percentage binding of the second NbHCV in the presence of a pre-bound first NbHCV, normalized to the binding of the second NbHCV in the absence of competition.

Next, NbHCV-NbHCV competition experiments were performed using biolayer interferometry (BLI) (Fig. 4C). Strep-tagged H77 E2c3 was immobilized on streptavidin biosensors, and the first NbHCV was loaded at a saturating concentration until equilibrium binding was reached. The sensors were then transferred to wells containing a second NbHCV at a saturating concentration, and the resulting spectral shift was recorded. Competition was quantified as the percentage of binding of the second NbHCV in the presence of the first NbHCV relative to its binding in the absence of the competitor. The results indicate that NbHCVs can be classified into two main groups. The first includes the broadly neutralizing NbHCVs from families 1, 8, and 13, and the low neutralizing NbHCV173 (family 40), whereas the second includes the low neutralizing NbHCVs from families 15 and 17. The first group can be further divided into two groups: NbHCVs from families 1 and 8 versus NbHCVs from families 13 and 40. To verify these findings, we incubated H77 E2c3 with NbHCV91 and NbHCV135, or with NbHCV128 and NbHCV135, at a 1:1:1 molar ratio and tested trinary complex formation via SEC. The results confirm that NbHCV135 does not compete with the binding of NbHCV91 or NbHCV131 (Supplementary Fig. 4B).

### HDX-MS mapping reveals three structurally distinct epitope groups

To achieve high-resolution mapping of NbHCV epitopes, we performed hydrogen–deuterium exchange mass spectrometry (HDX-MS) experiments. HDX-MS measures the rate at which backbone amide hydrogens exchange with deuterium from deuterated water (D_2_O), a process that is highly sensitive to local structural dynamics and solvent accessibility. We compared the deuterium uptake of apo H77 E2c3 with that of H77 E2c3 in complex with each member of the 12-NbHCV panel. The proteins were digested by a co-immobilized AnPep/pepsin protease column, followed by deglycosylation using a PNGase Rc column to remove E2 N-linked glycans. This approach yielded peptide fragments covering 98% of the E2c3 sequence (Supplementary Fig. 5A). Proteins were incubated in deuterated buffer for varying time points. Differential deuterium exchange was calculated by subtracting the exchange percentage of the E2c3-NbHCV complex from that of apo E2c3, where positive values indicate reduced solvent accessibility upon Nb binding. The maximum difference occurred at the first time point (20 s) and was used for subsequent analysis (Supplementary Fig. 5C).

Analysis of the HDX-MS data revealed that the NbHCVs can be classified into three distinct groups based on their binding epitopes, in agreement with the binding competition results (Fig. 5). Class 1 NbHCVs target the β-sandwich region, primarily engaging β5 and the loop connecting β6 and β7, which flanks the CD81-binding loop. Structural analysis of the E2-CD81 complex (PDB: 7MWX) shows that the β5 strand undergoes conformational rearrangement upon CD81 binding, providing a structural basis for the observed competition with CD81 and the neutralization activity of this group. Class 2 NbHCVs recognize epitopes that span the front layer, β5, and the CD81-binding loop, consistent with their strong competition with CD81- and AR3-targeting Abs and their broad neutralization potency. Notably, although HDX-MS and competition assays classified NbHCV173 within this group, it exhibited limited neutralization activity, suggesting that epitope engagement alone may be insufficient for effective neutralization. Class 3 NbHCVs are low-neutralizing Nbs that bind a short, linear epitope within the back layer (β11), overlapping with the AR2 antigenic region, as previously described ^35^.

**Fig.5.**
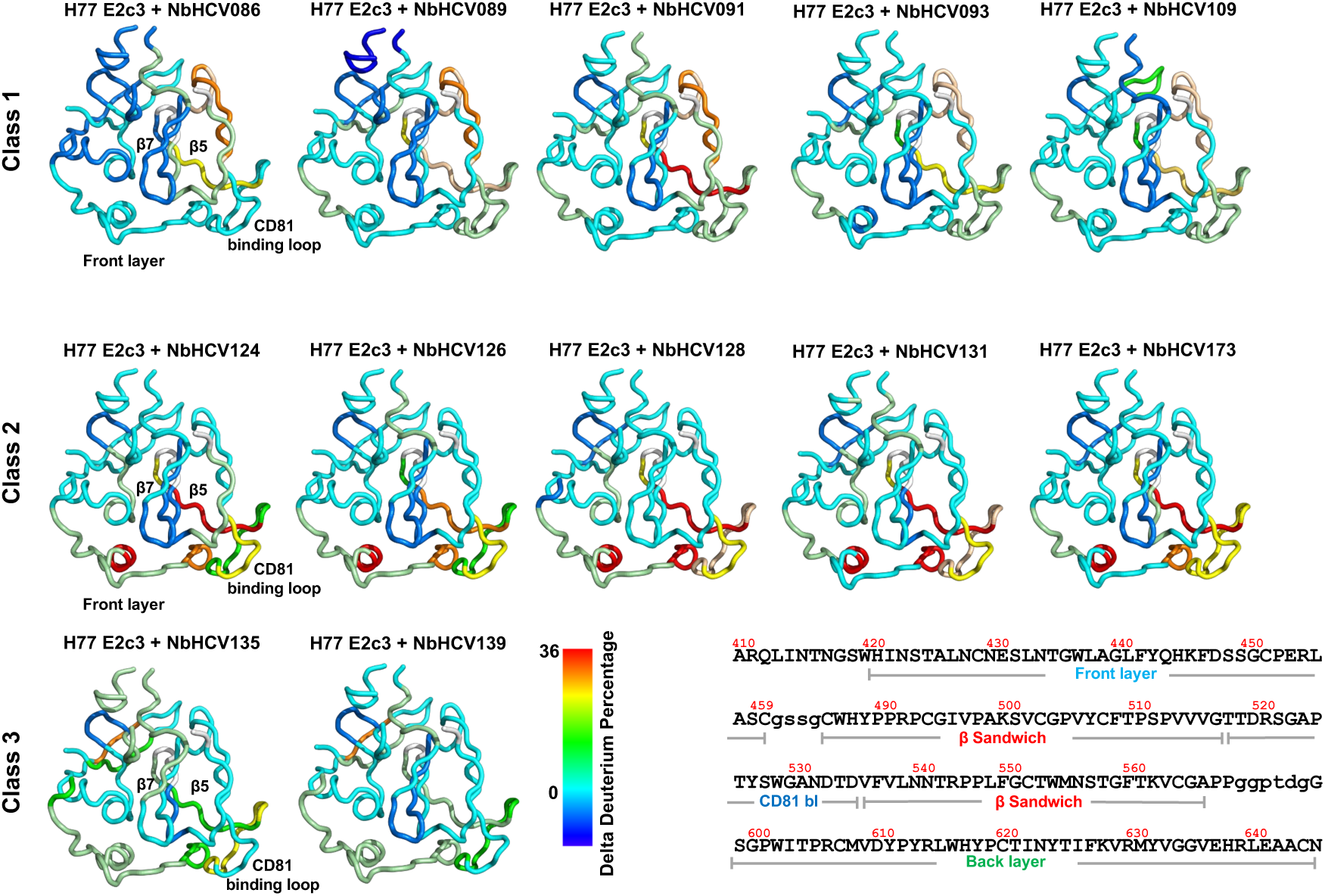
Epitope mapping of NbHCVs by HDX-MS. **(A)** Differences in deuterium uptake between E2c3-NbHCV complexes and unbound E2c3 after exposure of 20 seconds to D2O. Results are mapped onto the crystal structure of E2c3 with a color gradient indicating the percentage reduction in deuterium exchange for the 12-NbHCV panel. The percentage reduction is calculated as the decrease in deuterium uptake for each peptide in the E2c3–NbHCV complex relative to unbound E2c3. A high positive difference (red) highlights regions that are exposed in unbound E2c3 and protected upon NbHCV binding. Regions of E2 not covered by HDX peptides are shown in white.

A structural comparison of the putative epitopes of representative NbHCVs from each class with those of human and RM bnAbs (Table 1) revealed that class 2 NbHCV epitopes overlap with the E2 neutralization face and resemble the binding mode of the RM11-43 Abs class (Fig. 6A, B). In contrast, class 1 NbHCVs target an epitope outside the neutralization face, with a binding profile similar to that of hcab17. This region, corresponding to antigenic region 1 (AR1), was long considered a non-neutralizing site, targeted by the non-neutralizing Abs AR1A and E1 (Fig.6C) ^12,21,36^. However, recent work by Ogega et al. ^23^ demonstrated that AR1 can also be recognized by nAbs, e.g., hcab17, redefining it as a functionally relevant site of vulnerability outside the neutralization face. The ability of class 1 NbHCVs, which exhibit moderate neutralizing activity comparable to or exceeding that of hcab17, to engage this region further supports its role in neutralization and highlights its potential as a vaccine target. Collectively, these data expand the antigenic landscape of HCV E2 by identifying distinct epitope classes associated with neutralization. These insights establish new targets for the design of broadly protective HCV vaccines.

**Table 1.**
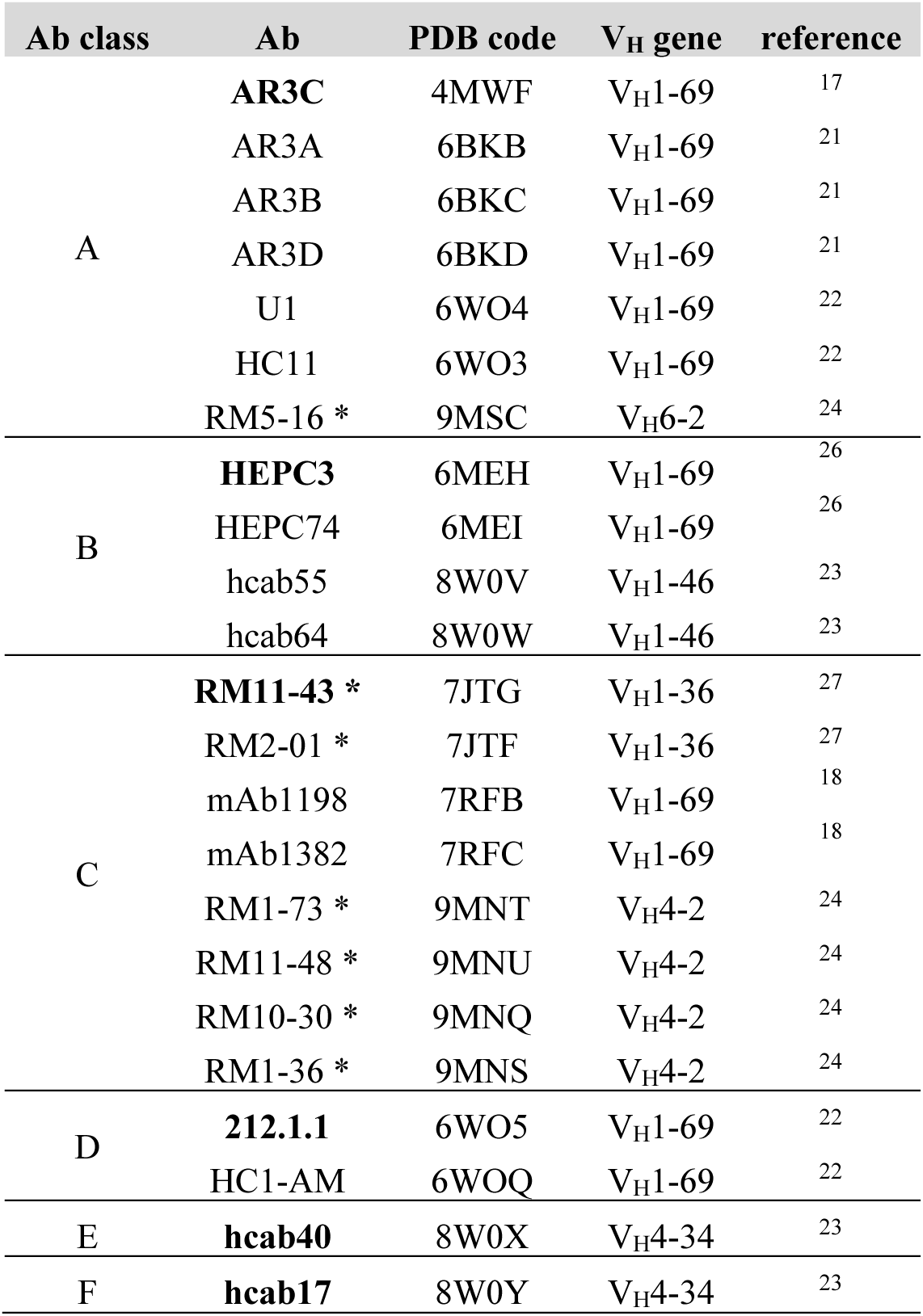
Classification of E2-specific bnAbs based on their binding epitopes. The representative Ab of each class (Fig. 6B) is marked in bold. RM Abs are marked with an asterisk.

**Fig.6.**
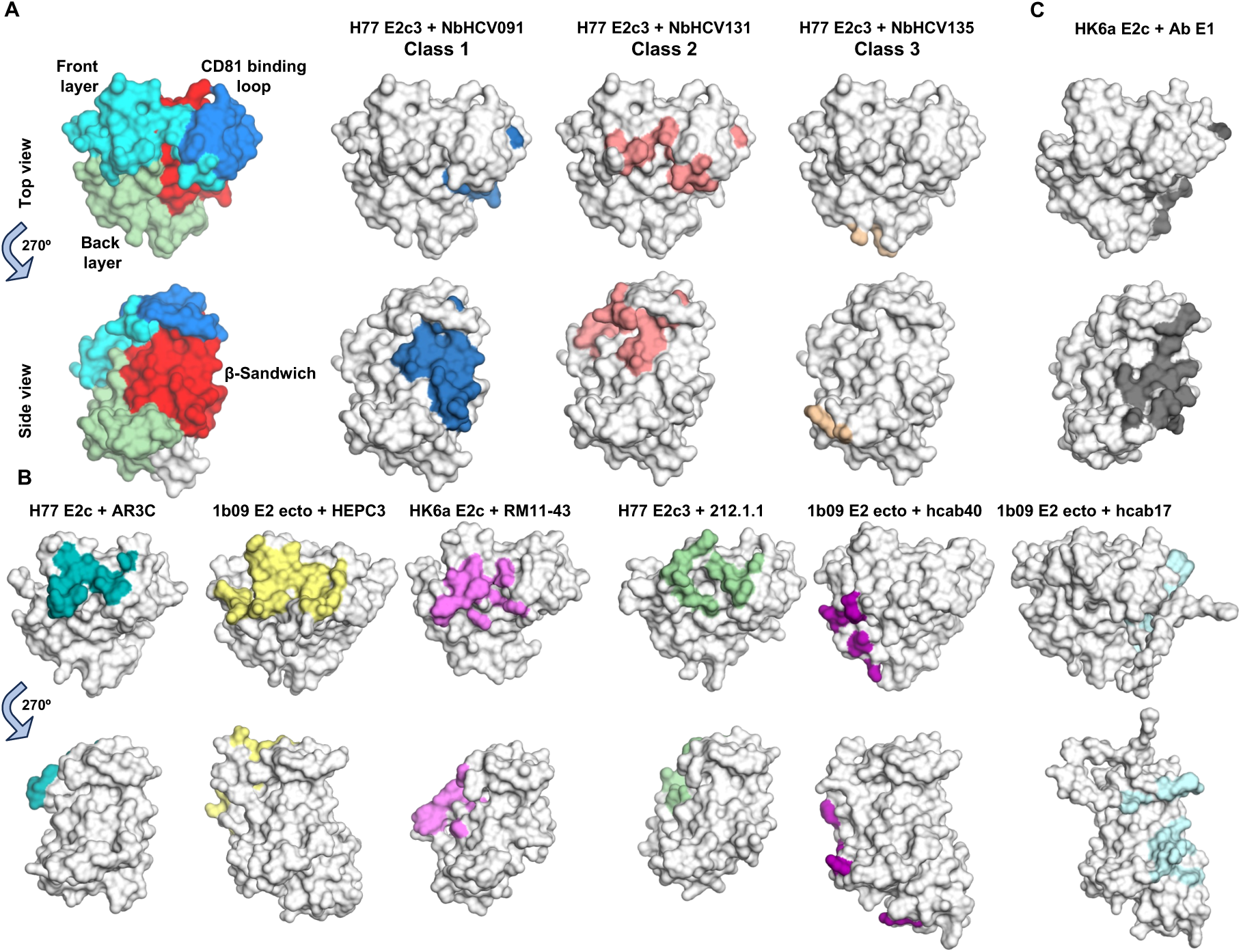
Comparison of NbHCV putative epitopes with epitopes of E2-specific bnAbs. **(A)** Putative epitopes of representative NbHCVs from each class, as defined by HDX analysis. E2c structures are shown in surface representation in both top and side views, with interacting residues highlighted. For reference, the structure of HK6a E2c3 is shown on the left, with the molecular surface colored by structural elements (see Fig.1A). **(B)** Epitopes of representative E2-specific bnAbs from distinct Ab classes. E2c structures are displayed in surface representation in top and side views, with epitope residues highlighted. **(C)** Epitope of the non-neutralizing Ab E1 that targets AR1.

## Discussion

The development of an effective HCV vaccine remains a global priority, particularly given ongoing transmission in high-risk populations and the epigenetic signature the virus leaves behind even after clearance. A major challenge in HCV vaccine design is the substantial genetic diversity and conformational flexibility of the viral envelope glycoproteins, underscoring the need for improved vaccines that elicit robust bnAb responses. In this study, we used the structural accessibility and diversity of camelid HCAbs to explore the antigenic landscape of the HCV E2 glycoprotein and identify previously unrecognized neutralization epitopes relevant to vaccine development.

HCAbs have been used to identify viral neutralization epitopes. Due to their small size, HCAbs can access conformationally flexible or hidden surfaces that may be poorly targeted by conventional Abs, thereby serving as valuable tools to guide improved antigen design. The human bnAb response against HCV appears to be dominated by V_H_1-69-encoded Abs that converge on overlapping epitopes within the E2 neutralization. While this focused targeting highlights the importance of the neutralization face, it may also limit the variety of antigenic sites that can be effectively targeted. Conversely, the diverse genetic architecture of camelid HCAbs broadens the range of possible paratope configurations, allowing alternative binding modes and revealing antigenic surfaces that are less targeted by typical human responses. By avoiding V_H_1-69 biases and immunodominance hierarchies, this approach offers a broader, less restricted view of E2 antigenicity, uncovering epitopes that could guide the design of immunogens with multiple neutralization sites to elicit more diverse and resilient nAb responses.

To maximize the diversity of elicited HCAbs, we immunized with both E2ΔTM and the truncated E2c3 core domain. We hypothesized that these immunogens would bias the antibody response toward distinct neutralization mechanisms. Specifically, E2ΔTM, which retains HVR1 and the stalk region, was expected to expose additional functional surfaces and potentially elicit Abs that interfere with viral entry through mechanisms other than direct CD81 blockade, such as by disrupting SR-BI engagement or inhibiting conformational rearrangements required for entry. In contrast, the structurally minimized E2c3 core, which prominently presents the conserved neutralization face, was anticipated to preferentially elicit CD81-blocking Abs. The immunization with these two antigens generated a highly diverse Ab repertoire, comprising 49 distinct families based on sequence conservation and phylogenetic clustering.

Immunization with E2c3 elicited a relatively weak Ab response, producing Nbs with low ELISA binding signals that were not further characterized. In contrast, E2ΔTM immunization generated a highly diverse Nb repertoire with strong cross-reactivity and broad neutralizing activity. Note that the E2c3 and E2ΔTM antigens were immunized in two different animals, which can also explain the difference in the immune response. Neutralization potency closely tracked with the ability to block CD81 engagement, underscoring the central role of receptor blockade in effective HCV neutralization.

Together, these findings suggest that, despite its structural simplification, E2c3 may lack critical conformational or contextual features necessary to elicit functional nAbs, whereas the more native-like architecture of E2ΔTM better preserves the antigenic determinants required for productive immune engagement.

Mapping of the NbHCV binding epitopes using competition and HDX experiments demonstrated that NbHCVs can be classified into three groups targeting distinct antigenic regions. Class 2 NbHCVs (except for NbHCV173) are the most potent, with high binding affinity and cross-neutralization potency, targeting the well characterized AR3. HDX experiments suggest that they resemble the binding mode of the RM bnAbs RM11-43 like Abs and the human bnAbs mAb1198 and mAb1382 (Fig.6 and table 1). These results are consistent with previous studies that demonstrated the neutralization face as the most potent and dominant neutralization site.

Nonetheless, a key objective of this study was to discover previously unrecognized neutralization epitopes. Antigenic region 1 (AR1), located in the back layer of the E2 core domain and mainly involving the β-sandwich, was long considered a non-neutralizing site. Abs targeting this region (e.g., AR1A and E1) typically showed little or no neutralization activity and were believed to recognize non-functional or transient conformations of E2. However, more recent research by Ogega et. al. ^23^ demonstrated that AR1 can also be targeted by nAbs such as the low neutralizer Ab hcab17. Consistent with these results, our data reveal that class 1 NbHCVs, which display moderate broad neutralization potency, target epitopes that overlap with the hcab17 binding site, further supporting the role of AR1 in the neutralization response against HCV.

Collectively, our findings expand the current understanding of the HCV E2 antigenic landscape by identifying both dominant and previously underappreciated neutralization sites. In particular, the ability of NbHCVs to uncover a functional role for AR1 highlights the importance of exploring beyond immunodominant epitopes. These insights provide a framework for the rational design of next-generation HCV immunogens that incorporate multiple vulnerable sites to elicit broader and more effective neutralizing Ab responses.

## Materials and methods

### Camelid immunization, construction of the NbHCV library, and production of a final HCV E2-specific Nb library

Immunization experiments were conducted in collaboration with the Nanobodies4Instruct Center in Belgium. To increase the chances of eliciting NbHCVs, two E2 antigens were used for camelid immunization: soluble H77 E2 ectodomain (H77 E2ΔTM, amino acids 384-645 of the H77 prototypic strain, UniProt number P27958) and soluble core domain (E2c3, E2 residues 412–645 with an internal truncation of VR2, VR3, and removal of the N448 and N576 glycosylation sites). Expression and purification of the antigens were performed as previously described ^21^. Camelid immunization, lymphocyte isolation, phage library preparation, and creation of the final panned Nbs library were carried out as previously detailed ^32^. In summary, two llamas (one per antigen) were immunized with highly purified E2 through six consecutive injections, and blood was collected five days after the final immunization. A serum binding assay indicated a high level of E2-specific HCAbs. Total RNA was extracted from peripheral blood lymphocytes, and cDNA was synthesized. The HCAbs variable regions were amplified and cloned into the pMESy4 vector to generate two E2-specific phage display libraries. Panning experiments were conducted on these libraries, and E2-specific clones were selected. The antigen-specific clones were sequenced to generate the final NbHCVs library.

### NbHCVs expression and purification

NbHCVs were expressed in Escherichia coli (E. coli) WK6 cells using the pMESy4 vector with a C-terminal His₆ tag and periplasmic secretion under the pelB leader ^32^. pMESy4 vectors were transformed and plated on Luria Broth (LB) media agar plates supplemented with ampicillin (Amp, 100 µg/ml) and 2% (wt/vol) glucose. For NbHCVs expression, a culture of Terrific Broth (TB) media supplemented with Amp, 0.1% (wt/vol) glucose, and 1 mM MgCl_2_ was grown until OD 600 = 0.6-0.7. Protein expression was induced by adding 1 mM of isopropyl-β-D-thiogalactopyranoside (IPTG), and the cells were grown overnight at 28°C.

On the following day, the bacterial cells were harvested by centrifugation at 9,000g for 15 minutes (min) at room temperature (RT). The cell pellet was resuspended in ice-cold TES buffer containing 200 mM Tris-HCl (pH 8.0), 0.6 mM EDTA, and 0.5 M sucrose (15 mL per liter of culture) and incubated for 1 hour at 4°C on an orbital shaker. Next, a double volume of TES/4 buffer was added to resuspend the pellet, and the cells were shaken for 45 min at 4°C. The suspension was centrifuged at 10,000g for 30 min at 4°C. The supernatant, Nb-containing fraction was recovered and loaded onto a His-Trap FF column (Cytiva) equilibrated with washing buffer containing 20mM Tris-HCl (pH 7.5) and 100mM NaCl. After loading, the column was washed with 10 column volumes (CV) of washing buffer, followed by 3-5 CV of washing buffer containing 10 mM imidazole. The Nbs were eluted from the column with 4-5 CV of elution buffer, which contained 20mM Tris pH 7.5, 200mM NaCl, and 250mM imidazole, and the protein-containing fractions were collected. SDS-PAGE verified protein expression. The eluted protein was concentrated using a 5 kDa cutoff concentrator (Vivaspin ®) and subjected to a second purification step on a Superdex 75 Increase 10/300 GL column (Cytiva) equilibrated with a buffer containing 20mM Tris-HCl (pH 7.5) and 150mM NaCl. The purified protein was concentrated, its final concentration was determined by absorbance at 280 nm, and it was stored at –80 °C.

### Expression of soluble E1E2 for ELISA binding experiments

The E1E2 genes (amino acids 192-745 based on the H77 sequence) of the following isolates were cloned into the pCMV vector for mammalian cell expression: H77 ^37^, HK6a ^38^, UKNP1.9.1 (Addgene 97374), UKNP1.10.1 (Addgene 97376), UKNP1.11.6 (Addgene 97382), UKNP1.16.3 (Addgene 98188), 1A72 (Addgene 177690), 1A138 (Addgene 177688), UKNP1.18.1 (Addgene 98190), 1B25 (Addgene 177691), 1B34 (Addgene 177692), 1B58 (Addgene 177693), UKNP3.1.2 (Addgene 98215), UKNP4.2.2 (Addgene 98268), and UKNP5.2.1 (Addgene 98385). HCV1 ^39^, HKP1a, CON1 ^40^, UKN1b12.6 ^41^, J6 ^42^, UKN3A1.28c 3a ^41^, S52 ^43^, ED43 ^44^, SA13 ^45^.

For E1E2 expression, Human Embryonic Kidney 293 cells (HEK 293T) were transfected using Polyethylenimine (PEI, Polysciences) at a 5:1, PEI:DNA, ratio. 72 hours post-transfection, cells were harvested with Trypsin (Biological Industries, Beit Ha’emek, Israel), pelleted by centrifugation at 400g for 5 min, and washed with cold PBS. The washed pellet was resuspended in cold lysis buffer containing 150 mM NaCl, 50 mM Tris (pH 7.5), 2 mM EDTA, and 0.5% Triton (100 μL of lysis buffer per 10^6^ cells) and incubated on ice for 30 min. The cell lysate was centrifuged at 20,000 × g for 15 min to remove cell debris, then aliquoted and stored at -80°C.

### Expression of Fc-CD81 LEL and domain-specific E2 Abs

The long extracellular loop of the tetraspanin CD81 (CD81 LEL; Uniprot number P6033, amino acids 113-202), fused to a C-terminal IgG Fc domain and a 6xHis-tag, was cloned into the phCMV1 vector (Fc-CD81-LEL). Fc-CD81-LEL was expressed in Expi 293F cells following the manufacturer’s instructions. The heavy chain (HC) and light chain (LC) sequences of AP33 ^34^, AR3A^21^, HC1AM ^22^, HEPC3 ^26^, and E1^12^ IgGs were cloned into the phCMV1 vector and expressed in Expi 293F or Expi CHO cells. Fc-CD81 and IgGs were purified using an HP-HiTrap^TM^ protein G column, followed by SEC chromatography with a Superdex 200 Increase 10/300 GL column in a buffer containing 100 mM NaCl and 20 mM Tris-HCl (pH 7.5).

### Expression and purification of H77 E2c3 and HK6a E2c3

Stable HEK293 GnTI^-^cell lines expressing the E2c3 core domain from HCV strains H77 and HK6a were maintained under standard culture conditions. E2c3 was purified using a custom AR3A-conjugated affinity column, followed by SEC chromatography with a Superdex 200 Increase 10/300 GL column in a buffer containing 100 mM NaCl and 20 mM Tris-HCl (pH 7.5), as previously described^21^.

### Enzyme-Linked Immunosorbent Assay (ELISA)

Lectin-capture ELISAs were performed to assess the binding of the NbHCVs to soluble HCV E2 (E2ΔTM or E2c3) and to E1E2 complexes from different HCV genotypes. For ELISA against E1E2, half-area 96-well plates (Greiner Bio-One, catalog number 675083) were pre-coated with Galanthus nivalis lectin (GNL; 5 µg/mL; Sigma-Aldrich) for 1 h at room temperature (RT) to enable capture of glycosylated E1E2 from cell lysates. For soluble E2, plates were coated with 5 µg/mL E2. Coated plates were washed with PBS containing 0.05% Tween-20 (PBST) and blocked with 5% (w/v) nonfat dry milk in PBS for 1 h at RT or overnight at 4°C. Following blocking, plates were washed and incubated with antigen, either soluble E2 (5 µg/mL) or E1E2-containing cell lysates, for 2 h at RT or overnight at 4°C. Plates were then washed and incubated with NbHCVs (5 µg/mL) for 1 h at RT. After washing, HRP-conjugated secondary antibodies were added for 1 h at RT (anti-His; ab1187, Abcam, 1:500 dilution, or anti-human IgG, 150217). Following incubation, the plates were washed, and TMB substrate (BioLegend) was added for color development. The reaction was stopped by adding 2 M sulfuric acid, and absorbance was measured at 450 nm using a microplate reader. All measurements were performed in duplicate. Background signal from antigen-free control wells was subtracted from all values, and mean absorbance values were calculated.

### Competition ELISA

For competition assays of NbHCVs with domain-specific monoclonal Abs (mAbs), E2-coated wells were first incubated with each mAb at saturating concentration (5 µg/mL) for 1 h at RT. Next, His-tagged NbHCVs were added at their EC_75_ concentration (Supplementary Fig. 4), plates were incubated for an additional 1 h, and NbHCVs binding was detected with an HRP-conjugated anti-His Ab. For competition assays of Nbs with the HEPC3 IgG and the CD81 receptor, captured E2 was first incubated with NbHCVs at a saturating concentration for 1 hour at RT, followed by the addition of HEPC3 IgG or Fc-CD81-LEL at its EC_75_ concentration (Supplementary Fig. 4) for an additional hour. Bound HEPC3 IgG or CD81 was detected using HRP-conjugated anti-human Fc antibody. Following incubation with the appropriate detecting antibody, plates were washed, developed with TMB, stopped with 2 M sulfuric acid, and absorbance was recorded at 450 nm. The percentage of inhibition was calculated using the following equation: % Inhibition = [1− (Signal with competitor/Signal with no competitor)]×100.

### HCV pseudoparticle production

HCV pseudoparticles (HCVpp) were produced as previously reported ^1^. In summary, HEK293T cells were co-transfected with the pNL4-3.Luc.R⁻E⁻ plasmid, which encodes HIV-1 Gag-Pol and a luciferase reporter gene (AIDS Reagent #3418), along with HCV E1E2 expression plasmids at a 1:4 ratio using PEI (5:1 PEI:DNA, w/w). Cells were maintained in DMEM supplemented with 10% FBS, sodium pyruvate, L-glutamine, and penicillin–streptomycin, and incubated at 37 °C with 5% CO_2_. After 72 h, supernatants containing HCVpps were collected, filtered through a 0.22 µm filter, and stored at 4°C for up to two weeks or at -80°C for long-term storage. For isolates that produced low infectivity signals (1A138, 1B25, UKNP1.9.1, UKNP1.10.1, UKNP4.2.2, and UKNP5.2.1), the HCVpps were concentrated 10-fold before neutralization assays to ensure comparable signal intensity across all preparations.

### Neutralization assay

HuH-7 cells were seeded in 96-well plates (1.5×10⁴ per well) and cultured overnight at 37°C and 5% CO_2_. HCVpp stocks were serially diluted and titrated on HuH-7 cells, and infectivity was quantified by measuring luciferase activity to determine the appropriate viral dilution for neutralization assays. For neutralization, HCVpps were mixed with NbHCVs at a concentration of 20 µg/ml or 2 µg/ml and incubated for 1 h at 37 °C before addition to HuH-7 cells. The NbHCVs concentration was selected to approximate the molar equivalent of conventional IgG-based assays, which are typically conducted at the range of 100-10 µg/ml, taking into account the substantially smaller molecular weight of nanobodies (∼15 kDa) compared to IgG (∼150 kDa) ^1,46^.

After 4-6 h, the medium was replaced with phenol-red-free DMEM, and the cells were incubated for 72 h. Cells were then lysed using a buffer that contained 25 mM glycylglycine, 15 mM MgSO₄, 4 mM EGTA, 5 mM NaOH (pH 7.8), 0.01% Triton X-100, and luciferase activity was measured using Bright-Glo™ (Promega). Relative light units (RLU) were normalized to mock controls, and percent neutralization was calculated relative to the signal of HCVpp-only infection.

### Biolayer interferometry (BLI)

Binding affinities were measured using BLI with Gator Bio. His-tagged Nbs (6.25 µg/ml in 20 mM Tris-HCl pH 7.5, 100 mM NaCl) were immobilized on anti-His biosensors. Serial two-fold dilutions of H77 or HK6a E2c3 (20–640 nM) were tested for association and dissociation kinetics. The equilibrium dissociation constant (K_D_) was calculated as k_Off_/k_On_.

NbHCV-NbHCV competition experiments were performed to assess whether pairs of nanobodies could bind simultaneously to E2c3 (H77). Strep-tagged E2c3 was immobilized onto Streptavidin (SA) biosensors (Gator Bio) at a loading concentration of 10μg/ml. After immobilization and baseline stabilization, sensors were dipped into wells containing the first NbHCV at 7.5μg/ml, a saturating concentration that ensured maximal binding. Once equilibrium binding of the first NbHCV was achieved, sensors were transferred to wells containing the second NbHCV at 7.5 μg/ml, and association was recorded in real time.

### Hydrogen-Deuterium Exchange Mass Spectrometry (HDX-MS)

The E2c3-NbHCV complexes were formed by incubation of purified H77 E2c3 and each NbHCV in a molar ratio of 1:1.1 (E2:NbHCV) at RT for 1 hour, followed by SEC (Superdex 200) to remove unbound NbHCV using 20 mM tris and 100 mM NaCl (pH 7.5) buffer. Next, the complexes were subjected to Hydrogen-Deuterium Exchange Mass Spectrometry (HDX-MS) to identify epitope protection patterns. For peptide mapping, 100 pmol of H77 E2c3 was mixed in a 1:1 volume ratio with 1 M glycine, 2 M urea, and 200 mM TCEP, pH 2.3, and then injected into a co-immobilized AnPep/pepsin protease column (AffiPro, Czech Republic) connected to a PNGase Rc glycanase column (AffiPro, Czech Republic). An automatic HDX workstation was used for sample handling and injection. The generated peptides were trapped and desalted with a Microtrap column (Luna Omega 5 μm Polar C18, 100 Å Micro Trap 20 × 0.3 mm) for 3 min at a flow rate of 0.2 ml/min, using an isocratic pump delivering 0.4% aqueous solution of formic acid. Both enzyme columns and the trap column were cooled to 4°C. After 3 min, peptides were separated on a C18 reversed-phase column (Luna® Omega 1.6 μm Polar C18, 100 Å, 100 × 1.0 mm) with a linear gradient of 5-45% solvent B over 8 minutes. Solvent A consisted of 2% acetonitrile and 0.4% formic acid in water, while solvent B was 95% acetonitrile, 5% water, and 0.4% formic acid. The analytical column was maintained at 4°C. Peptides were detected using a TimsToF Pro mass spectrometer (Bruker Daltonics) operating in parallel accumulation and serial fragmentation mode. Data were processed with DataAnalysis 5.1 software (Bruker Daltonics). Peptides were identified using Mascot against a database containing the H77 E2c3 sequence.

The HDX was initiated by a 10-fold dilution of H77E2c3 and E2c3-NbHCV complexes in a deuterated buffer (20 mM Tris pD 7.5, 100 mM NaCl). After 20 seconds, 2 min, 20 min, and 2 h of incubation in deuterated buffer, the reaction was quenched by adding 1 M glycine, 2 M urea, and 200 mM TCEP at pH 2.3. The 20-second and 20-minute time points were analyzed in triplicate. Sample handling was performed by an automated HDX workstation. Peptides were separated using a linear gradient of 5-45% solvent B over 8 minutes, where solvent A consisted of 2% acetonitrile and 0.4% formic acid in water, and solvent B was 95% acetonitrile with 5% water and 0.4% formic acid. The mass spectrometer (timsToF Pro) was operated in positive MS mode. Spectra of partially deuterated peptides were processed with Data Analysis 5.1 (Bruker Daltonics, Billerica, MA) and with in-house software DeutEx.

## Acknowledgements

This work is funded in part by the Israeli Science Foundation (ISF), grants #1600/21 and #1818/21 (to N.T.) and by the United States-Israel Bi-national Science Foundation (BSF), grant #2021-165 (to N.T. and M.L.). We acknowledge the support and use of resources of Instruct-ERIC (PID8041), part of the European Strategy Forum on Research Infrastructures (ESFRI), and the Research Foundation - Flanders (FWO) for their support of the Nanobody discovery.

We acknowledge the Structural mass spectrometry core facility in BioCev - project Instruct-ERIC (PID20617), part of the European Strategy Forum on Research Infrastructures (ESFRI), and the Research Foundation - Flanders (FWO) for their support of the mass spectrometry (HDX experiments).

H.T., R.F., and N.T. designed the project; E.G. expressed the antigen for immunization; E.P. and N.B. performed the immunization experiments and discovered E2-specific NbHCVs; H.T. and J.W. expressed and purified the NbHCVs and carried out ELISA binding experiments; H.T. and I.Y. expressed the E2c3 antigens; H.T. conducted BLI binding experiments, competition assays, and neutralization experiments; P.P. performed HDX experiments. H.T., R.F., P.P., M.L., and N.T. analyzed results and wrote the manuscript with support from all authors. The authors declare no competing interests. All data used to understand and evaluate the conclusions of this research are available in the main text and supplementary materials. Additional data related to this paper may be requested from the authors.

---

**Supplementary Fig.1.**
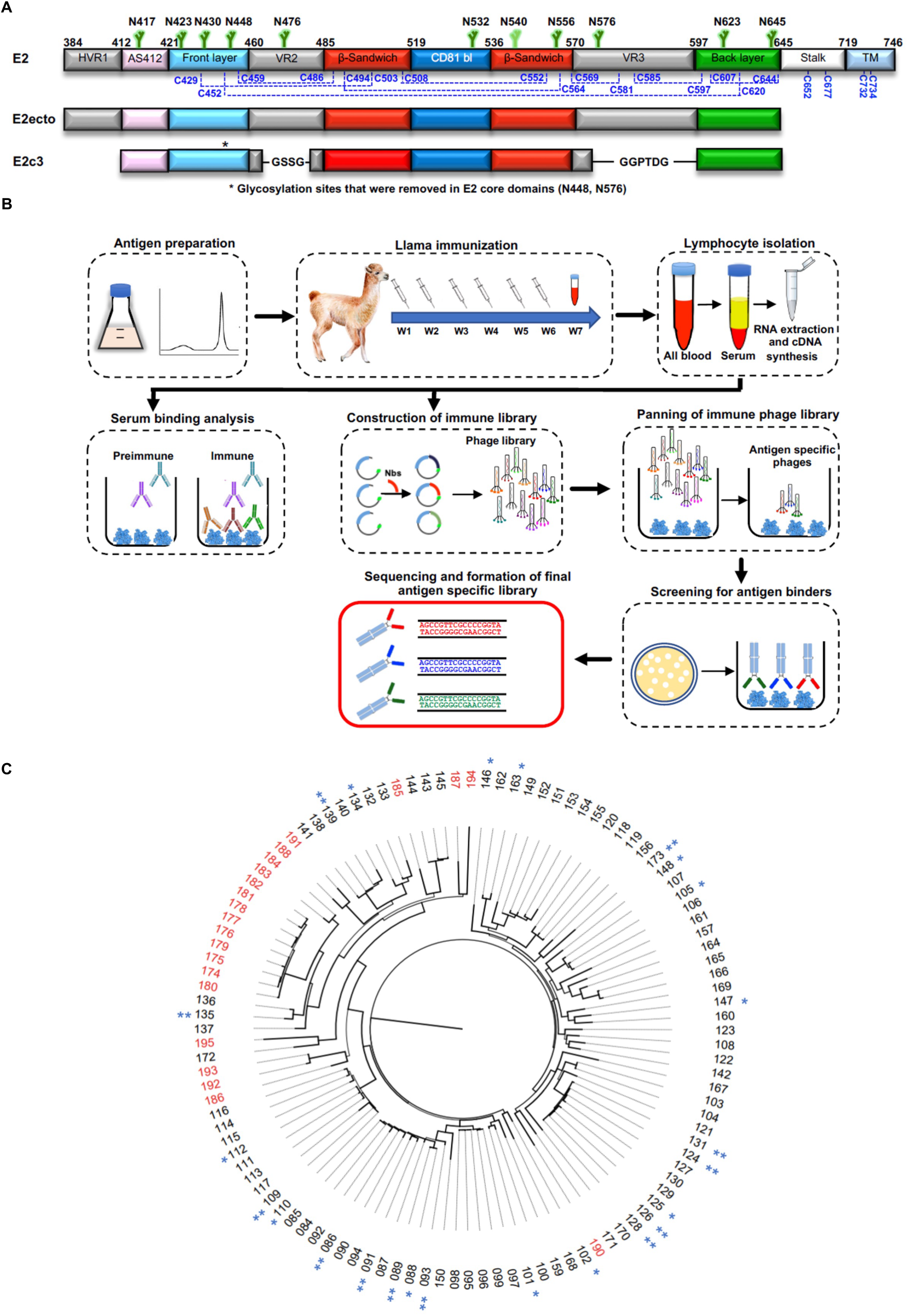
Generation and characterization of an E2-specific NbHCV library. **(A**) Schematic representation of the HCV E2 Env protein, E2 ectodomain, and E2 core domain colored by structural component. Numbering is based on the H77 prototypic strain. N-linked glycans are shown in green, and the conserved cysteines are in blue. For E2, disulfide bonds, based on the H77 E2C-AR3C structure, are indicated with blue dashed lines. HVR1, hypervariable region 1; AS412, antigenic site 412; VR, variable region; CD81 bl, CD81 binding loop; TM, transmembrane. **(B**) Schematic summary of the workflow used to generate the antigen-specific NbHCV libraries. **(C**) Maximum-Likelihood Phylogenetic trees were inferred from an amino acid alignment using FastTree v2.1.11, using the LG substitution model with a gamma distributed rate heterogeneity across sites. NbHCVs elicited by immunization with E2ΔTM are shown in black, whereas NbHCVs elicited by E2c3 are shown in red. NbHCVs included in the 12-NbHCV panel are indicated by double asterisks (**), while those included in the 24-NbHCV panel but not in the 12-NbHCV panel are indicated by a single asterisk (*).

**Supplementary Fig.2.**
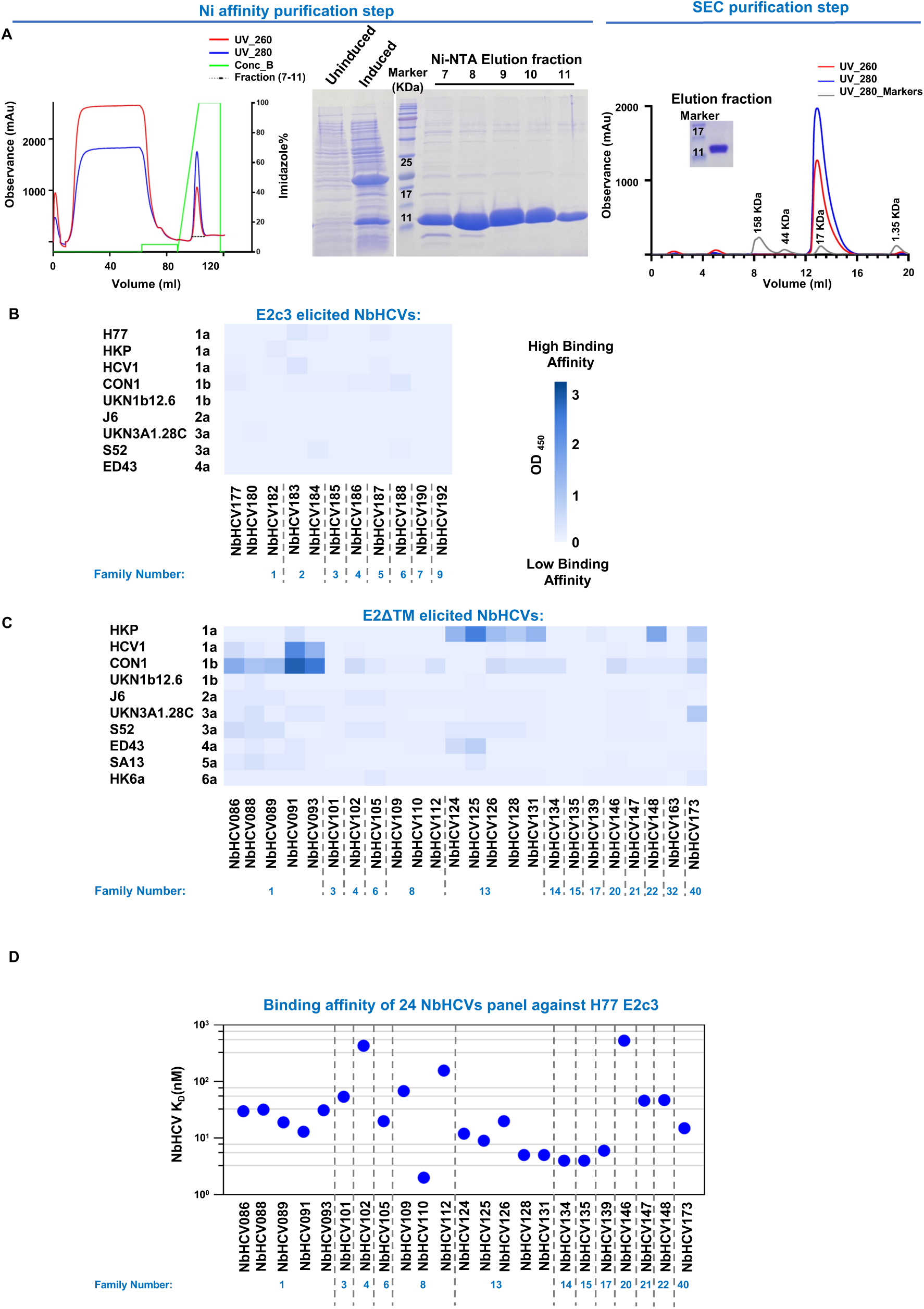
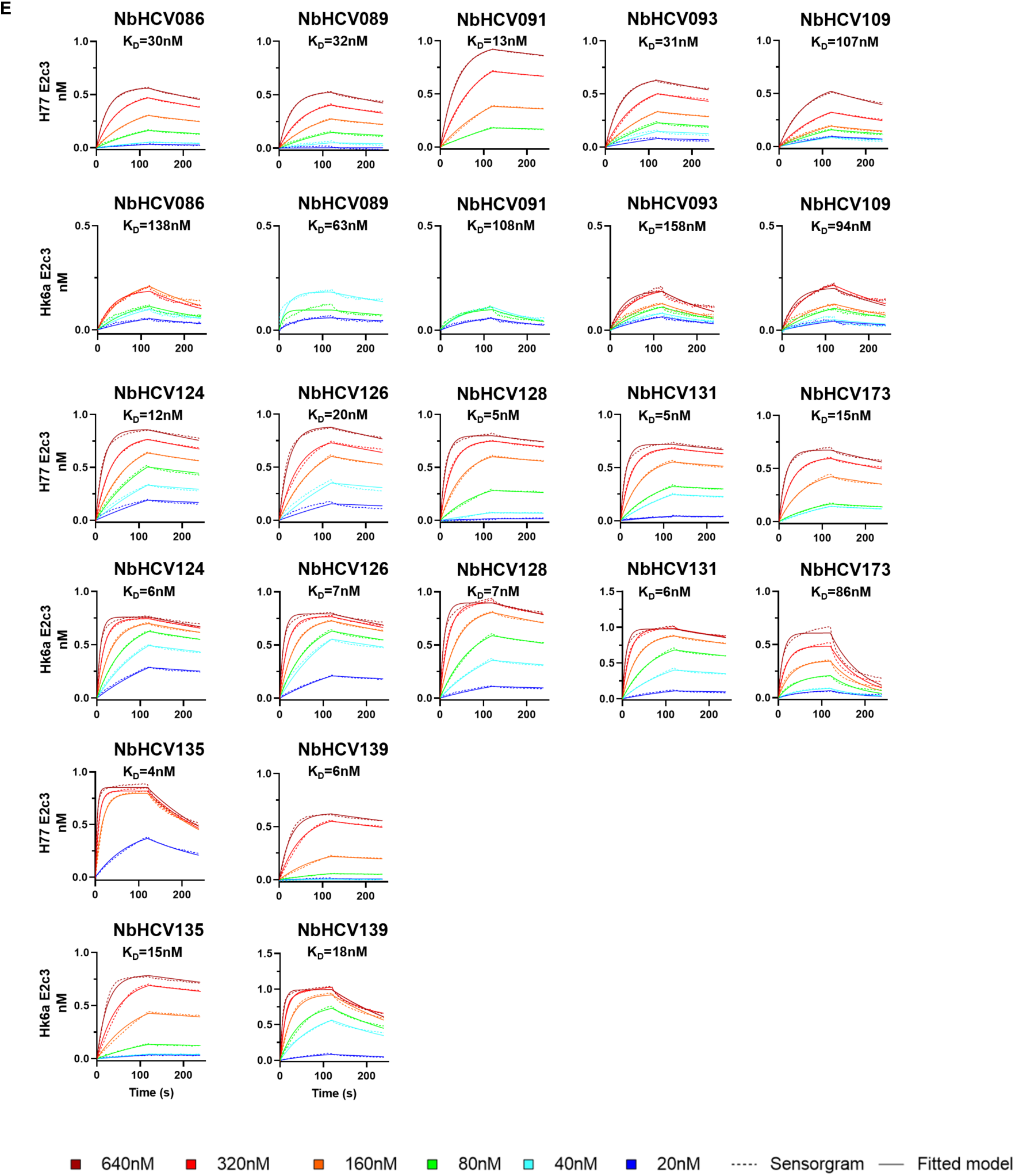
High-affinity cross-binding of NbHCVs to HCV envelope antigens across genotypes. **(A)** Purification of the NbHCVs. The purification results for NbHCV094 as a representative example of the NbHCV purification process. The NbHCVs were purified using a His-Trap FF affinity column (left). Protein expression was verified by SDS-PAGE (middle). Elution fractions were analyzed by a Superdex 75 size-exclusion column, which indicated the purified monomeric fraction of the NbHCV (right). (**B and C**) ELISA binding analysis of representative NbHCVs elicited by E2c3 immunization (**B**) or the 24-panel from E2ΔTM immunization (**C**) against a panel of native E1E2 HCV isolates from Gt1-Gt4. (**D**) Binding affinity determination of the 24 NbHCV panel against H77 E2c3 antigen. K_D_ values were determined by BLI. (**E**) BLI analysis of the 12-panel binding to soluble H77 and HK6a E2c3s. Sensorgrams were obtained using six titration series (starting with 640). K_D_ values were calculated from a 1:1 global fitting model (solid line) and are indicated next to the sensorgrams.

**Supplementary Fig.3.**
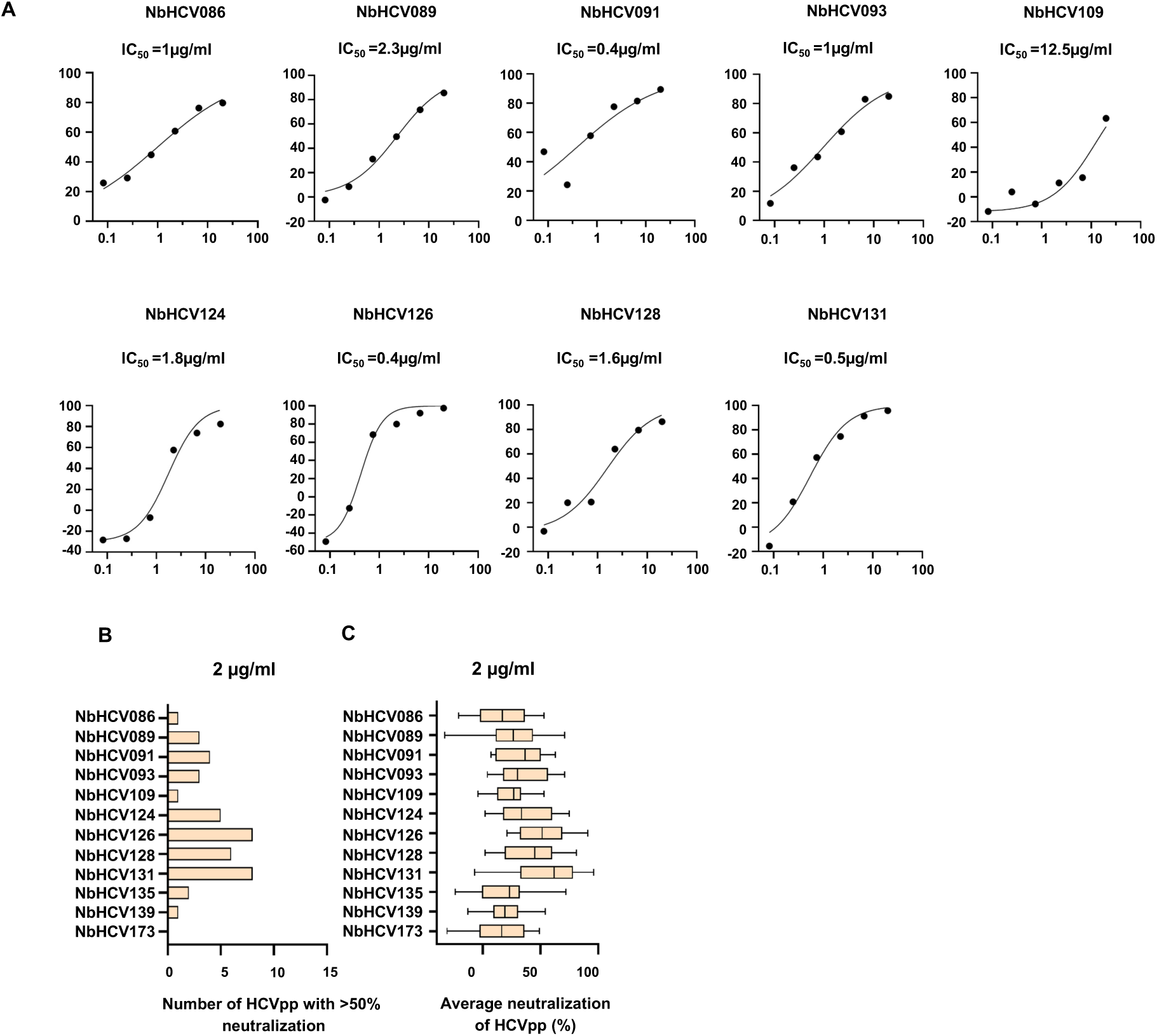
Cross-neutralization ability of the NbHCVs. (**A**) Dose response neutralization curves showing the percentage of neutralization by the 12-NbHCV panel against H77 HCV pseudoparticles (HCVpp). Half-maximal inhibitory concentration (IC_50_) values were calculated using nonlinear regression with a four-parameter logistic model in Prism. NbHCV135, NbHCV139, and NbHCV173 exhibited weak neutralization activity and did not achieve 50% neutralization. Therefore, IC_50_ values could not be determined. (**B**) The number of HCVpp with more than 50% neutralization for each NbHCV at a concentration of 2 µg/ml. (**C**) The average neutralization of each NbHCV against the HCVpp panel at 2 µg/ml. Lines, boxes, and whiskers represent medians, 25%–75%, and 5%–95% percentiles, respectively.

**Supplementary Fig.4.**
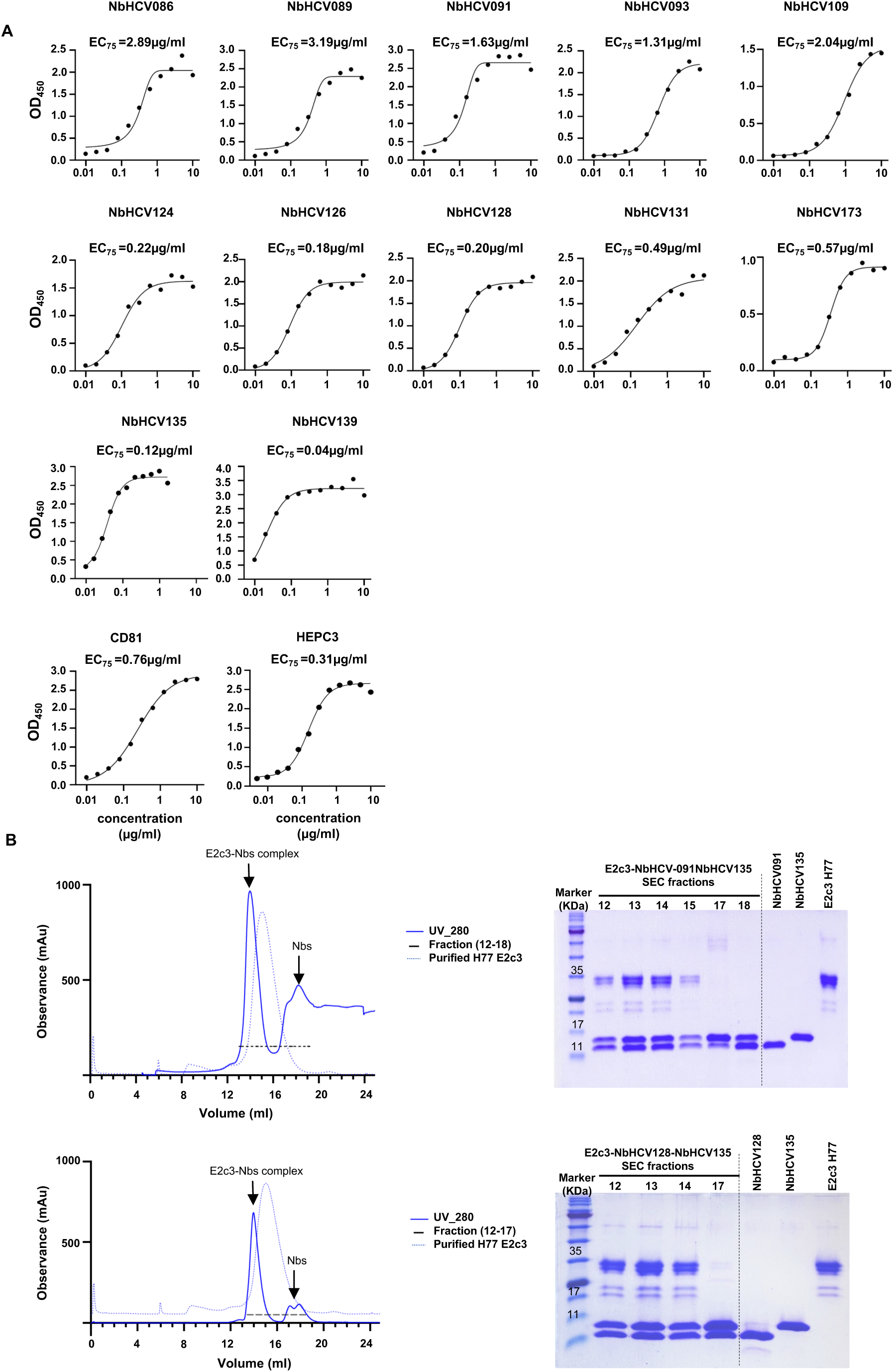
Epitope mapping of NbHCVs. **(A)** Dose response ELISA binding curves showing binding of the 14-NbHCV panel to H77 E1E2, measured as optical density at 450 nm (OD_450_). EC_75_ values were calculated by nonlinear regression using a four-parameter logistic model in OriginPro. **(B)** H77 E2v3-NbHCV091-NbHCV135 and H77 E2v3-NbHCV128-NbHCV135 complexes were formed by incubating purified H77 E2c3 with the NbHCVs at a molar ratio of 1:1.1:1.1 (E2:NbHCV:NbHCV), followed by SEC to test the competition between NbHCV091 and NbHCV128 for NbHCV135. The SEC results show that trinary complex formation is demonstrated on the left. The formation of these complexes was confirmed by SDS-PAGE, shown on the right.

**Supplementary Fig.5.**
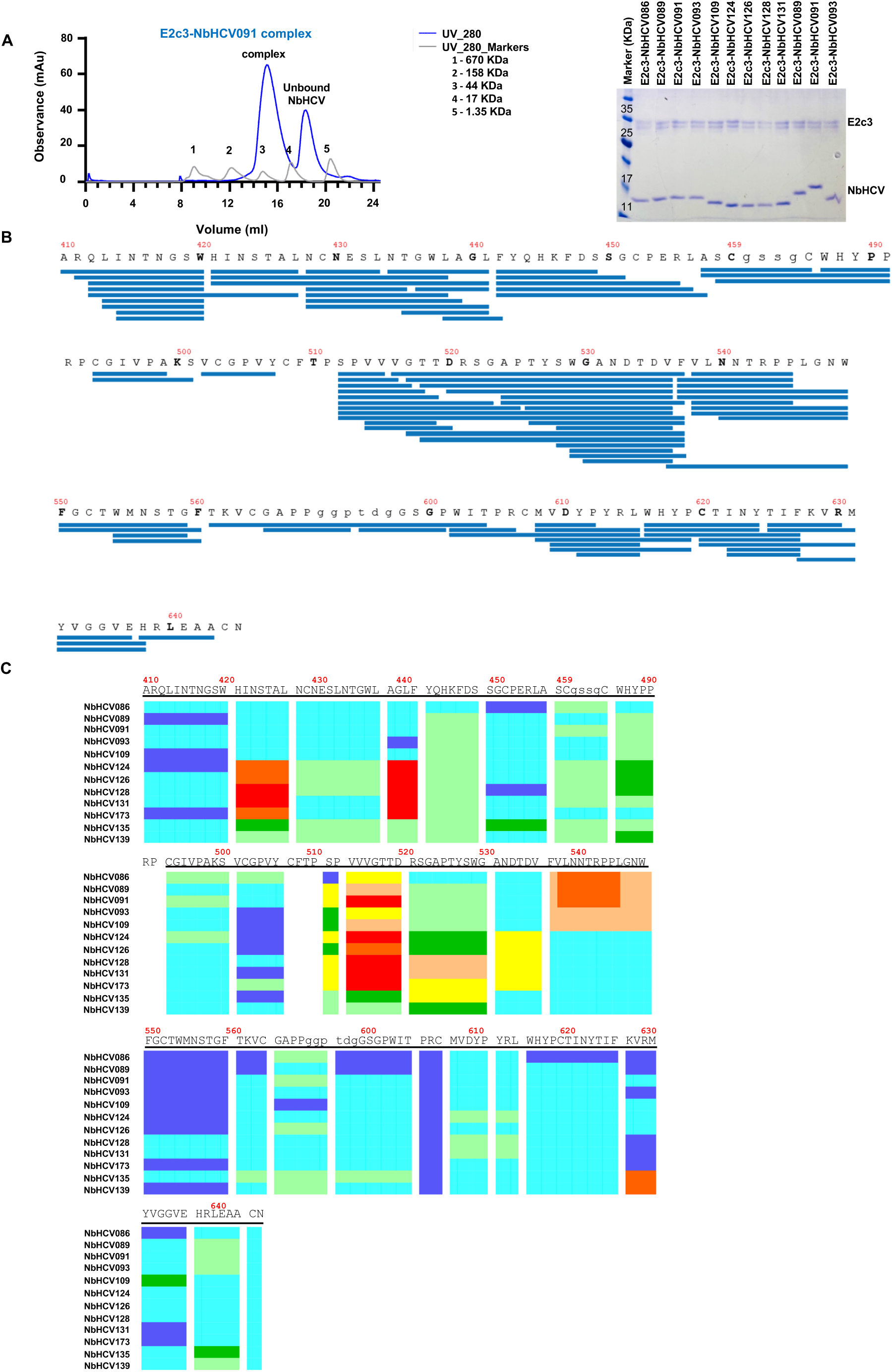
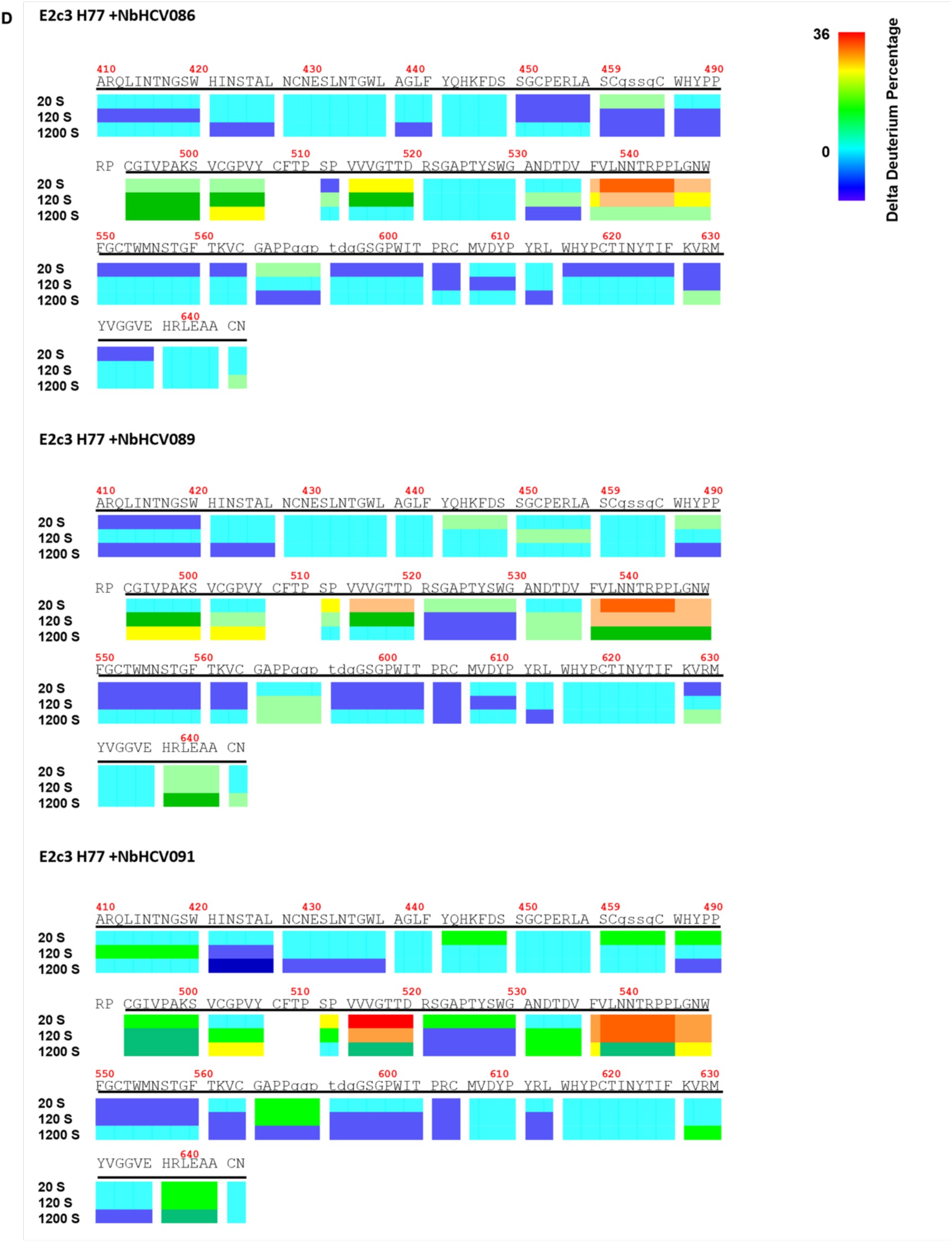
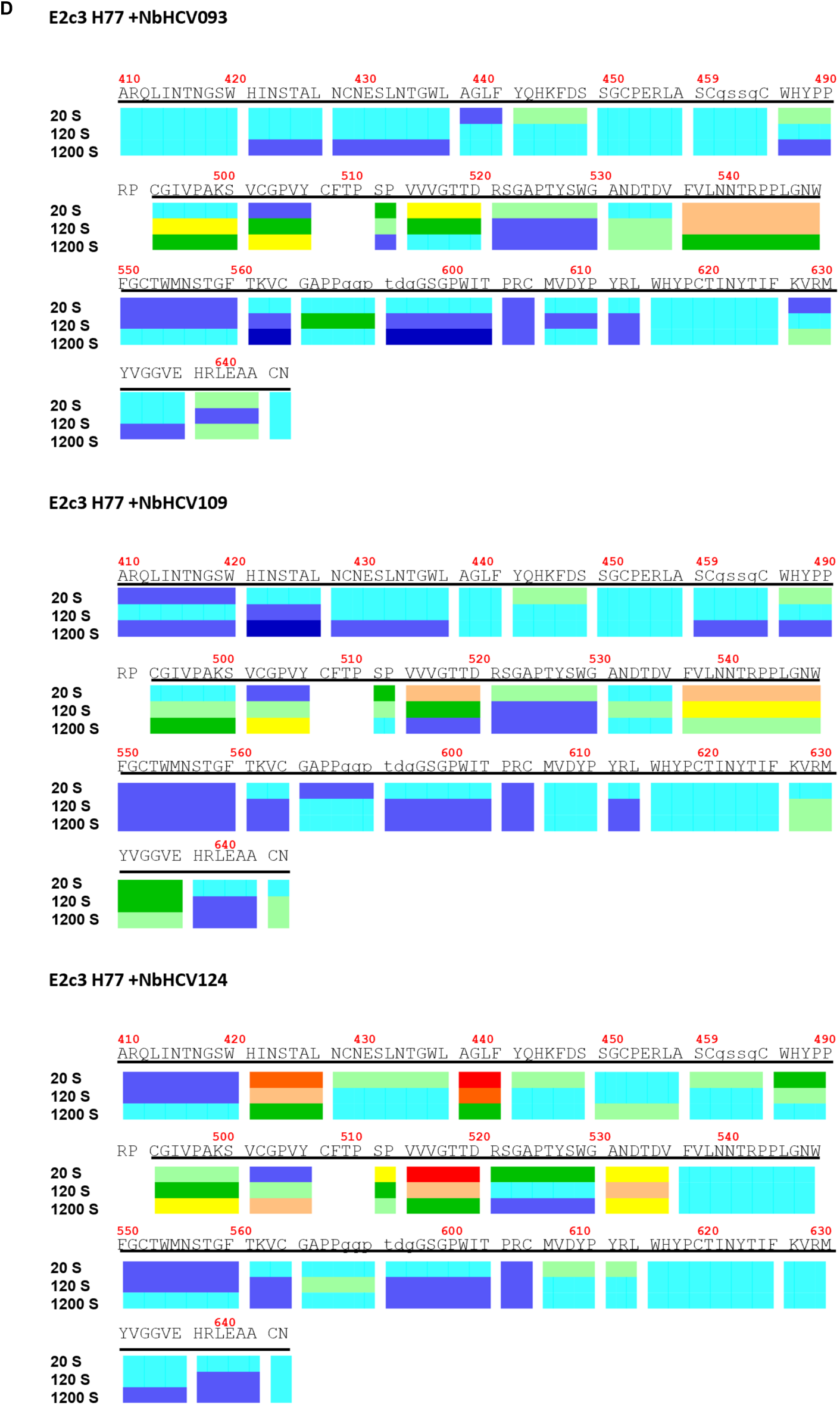
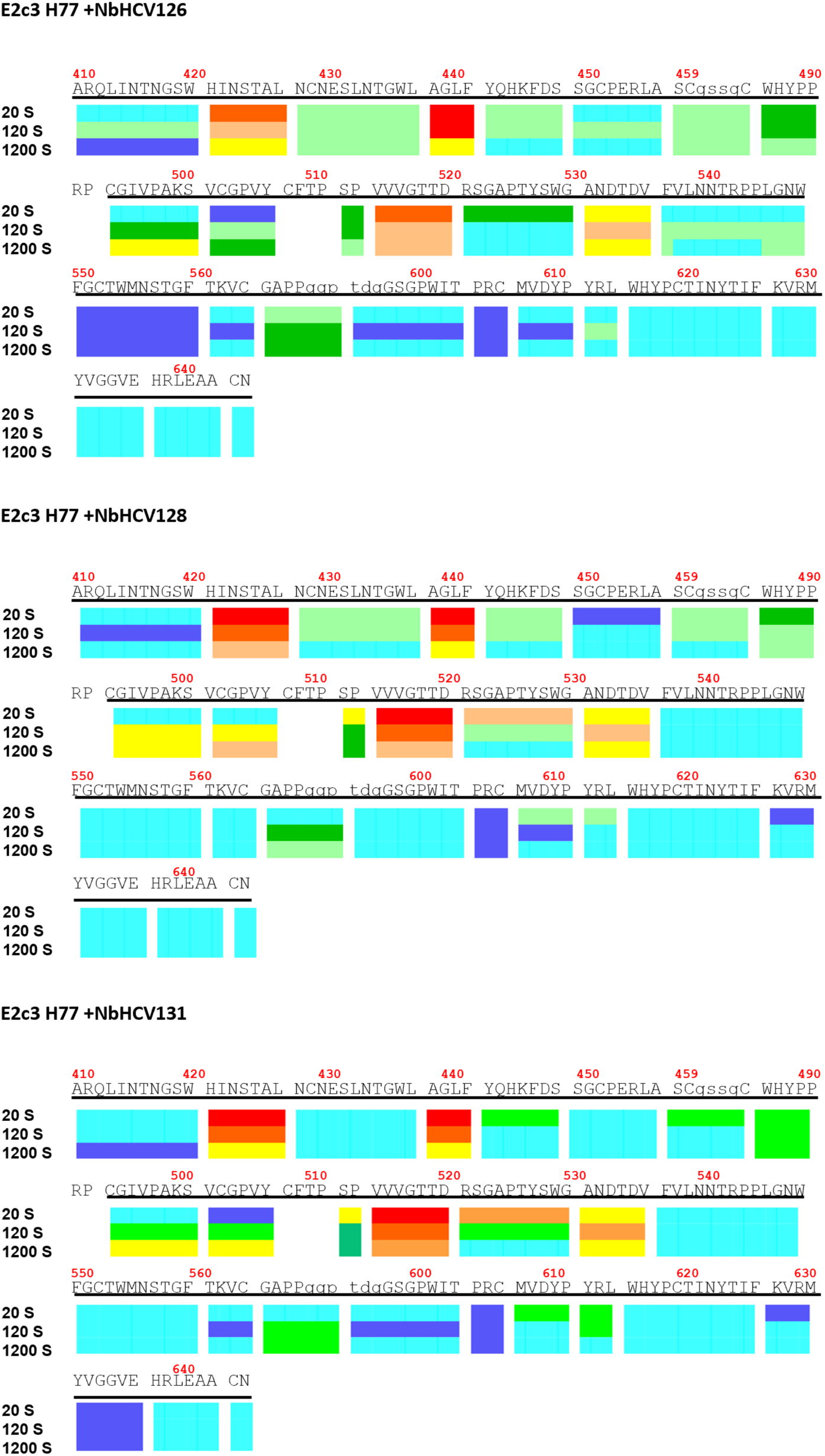
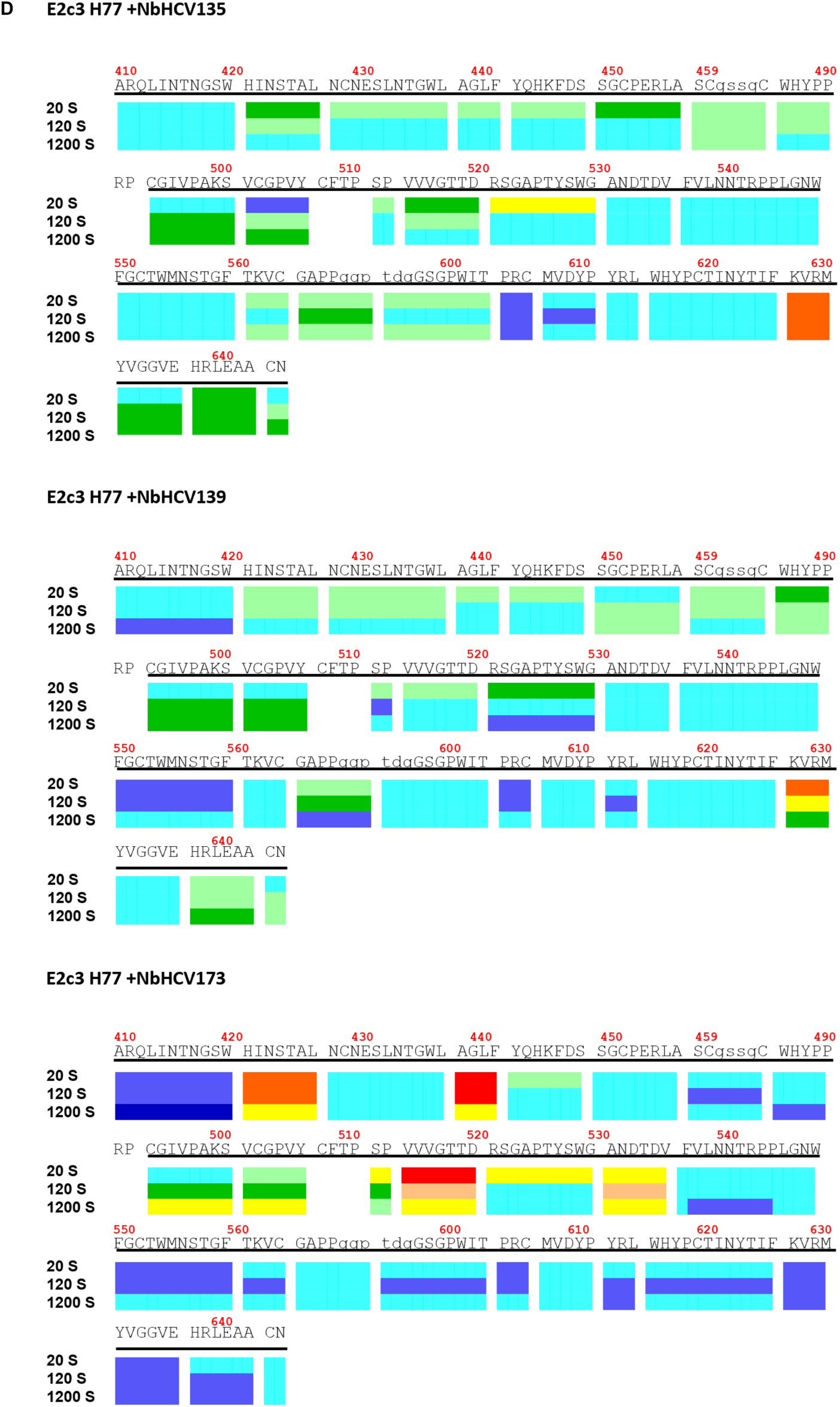
Epitope mapping of NbHCVs by HDX-MS. **(A)** H77 E2v3-NbHCV complexes for HDX experiments. H77 E2v3-NbHCV complexes were formed by incubating purified H77 E2c3 with each NbHCV from the 12-panel set at a molar ratio of 1:1.1 (E2:NbHCV), followed by SEC to remove unbound NbHCV. The SEC result for the E2v3-NbHCV091, a representative example, is shown on the left. The formation of all complexes was confirmed by SDS-PAGE, shown on the right. **(B)** HDX-MS peptide coverage of H77 E2c3. Peptides identified by mass spectrometry after HDX analysis of E2c3-NbHCV complexes are mapped onto the E2c3 amino acid sequence. (**C** and **D**) Differences in deuterium uptake between E2c3-NbHCV complexes and unbound E2c3 after exposure to D_2_O for different durations. In (**C**), results for all complexes at the 20 s time point are shown, while in (**D**), results for each complex are displayed across multiple time points (20s, 120s, and 1200s). Results are shown as a heat map indicating the percentage reduction in deuterium exchange for the 12-NbHCV panel. The percentage reduction is calculated as the decrease in deuterium uptake for each peptide in the E2c3– NbHCV complex relative to unbound E2c3. A high positive difference (red) highlights regions that are exposed in unbound E2c3 and protected upon NbHCV binding. Regions of E2 not covered by HDX peptides are shown in white.

**Supplementary Table 1. The NbHCV antibody library.** List of NbHCV Nb isolated following immunization with the H77 E2DelTM and H77 E2c3 antigens, grouped into sequence families. The lengths of the CDRs are indicated. NbHCV Nb included in the 53-, 24-, and 12-member panels are marked.

